# Oncogenic ESR1 fusions form chromatin-anchored condensates that remodel nuclear architecture

**DOI:** 10.64898/2026.10.01.755872

**Authors:** Ana-Maria Gherghelas, Natalia Fili, Adam Boulger, Alison E. Twelvetrees, Lin Wang, Lewis Frame, Jamieson A.L. Howard, Matthew D. Newton, Mark C. Leake, Christopher P. Toseland

## Abstract

Estrogen receptor (*ESR1*) fusions are recurrent drivers of endocrine-resistant breast cancer, yet their mechanisms of action remain poorly understood. Here, we show that *ESR1-e6* fusion proteins, including *ESR1-DAB2*, *ESR1-SOX9*, and *ESR1-YAP1*, assemble into nuclear condensates across multiple cell types. Sequence analysis reveals that these fusions combine chromatin-targeting domains with increased intrinsic disorder, promoting multivalent interactions and phase separation. Live-cell imaging, fluorescence recovery after photobleaching, and mechanical perturbation demonstrate that condensates exhibit liquid-like properties, but progressively mature into stable, viscoelastic assemblies constrained by chromatin. Disruption of DNA binding abolishes condensate formation, indicating that chromatin engagement is essential for their assembly and organisation. *ESR1* fusion condensates localise to chromatin-depleted regions, reduce nuclear stiffness, and promote chromatin decompaction, consistent with active remodelling of nuclear architecture. We propose that *ESR1* fusions exploit phase separation to generate chromatin-anchored assemblies that reorganise the nuclear microenvironment, providing a physical framework through which *ESR1* fusions may support sustained transcriptional activity, endocrine resistance and tumour progression.

## INTRODUCTION

Breast cancer remains the most diagnosed malignancy worldwide and the leading cause of cancer-related mortality among women (1, 2). Approximately 70% of breast cancers are estrogen receptor-positive (ER+), characterised by the expression of estrogen receptor alpha (ERα), a ligand-activated transcription factor encoded by *ESR1* (2, 3). ERα plays a central role in regulating genes involved in proliferation, survival, and differentiation through genomic and non-genomic mechanisms. In the canonical genomic pathway, ligand binding induces receptor dimerisation, nuclear translocation, and binding to estrogen response elements (EREs), enabling recruitment of co-regulators to control gene expression (4, 5). Disruption of this tightly regulated process leads to constitutive downstream signalling and activation of transcriptional programmes that promote tumour progression.

Although endocrine therapies, including aromatase inhibitors, selective ER modulators (SERMs), and selective ER degraders (SERDs), have substantially improved clinical outcomes, therapy resistance frequently emerges in a significant proportion of advanced cases (3, 6). This is often driven by mutations within the ligand-binding domain (LBD) – the primary therapeutic target – which confer ligand-independent activity or hypersensitivity to estrogen (7, 8).

More recently, *ESR1* gene fusions have emerged as an alternative mechanism of resistance. These rearrangements typically retain the DNA-binding domain (DBD), while eliminating the LBD, generating chimeric proteins with constitutive, hormone-independent activity that are associated with aggressive, therapy-resistant disease (9–13). Despite their growing clinical relevance, the molecular mechanisms underlying their activity remain poorly defined.

Beyond constitutive activation, transcription factor fusions can acquire biophysical properties that reshape gene regulation. Liquid–liquid phase separation (LLPS) has been recognised as a fundamental process that organises transcription within the nucleus, whereby multivalent interactions drive the formation of biomolecular condensates that concentrate transcriptional machinery (14, 15). Increasing evidence shows that oncogenic fusion proteins exploit phase separation to reorganise chromatin architecture and sustain aberrant transcriptional programmes (16, 17). *ESR1* fusion proteins retain DNA-binding capacity while incorporating intrinsically disordered regions from their partners, raising the possibility that these structural features promote phase separation and condensate formation. However, the contribution of condensates to *ESR1* fusion-driven phenotypes remains unexplored.

To address this, we investigated the nuclear organisation of recurrent patient-derived *ESR1* fusion proteins, focusing on *ESR1–DAB2*, *ESR1–SOX9* and *ESR1–YAP1*. Using complementary imaging and biophysical approaches, we show that *ESR1* exon 6 fusions form nuclear condensates across multiple cellular contexts, indicating that higher-order assembly is an intrinsic consequence of fusion architecture. We further reveal that these condensates display liquid-like properties, but progressively mature into stable, chromatin-constrained assemblies.

Together, our findings support a model in which *ESR1* fusions couple chromatin-binding capacity with partner-derived disorder to generate phase-separated nuclear assemblies. These condensates are not passive accumulations of fusion protein, but chromatin-anchored structures capable of reshaping the local nuclear environment. Our findings provide a mechanistic framework linking *ESR1* fusion architecture to altered chromatin organisation, hormone-independent transcriptional activity and endocrine-resistant breast cancer. Defining the formation, maturation and chromatin interactions of *ESR1* fusion condensates may therefore reveal new biomarkers and therapeutic vulnerabilities, while establishing phase separation as a central feature of *ESR1* oncofusion biology.

## MATERIALS AND METHODS

### Constructs

Plasmid constructs p.Halo-*ESR1*-e6, p.Halo-DAB2, p.Halo-*ESR1*-e6>DAB2, p.Halo-*ESR1*-e6>SOX9, p.Halo-*ESR1*-e6>NDP52, p.Halo-*ESR1*-e6>YAP1, p.Halo-*ESR1*-e7>POLH, p.Halo-*ESR1*-e6>ΔDBD>DAB2 (GeneArt ThermoFisher Scientific), cloned into the pcDNA 3.1(+) vector (ThermoFisher Scientific), were used in this study. Constructs were amplified in *Escherichia coli* bacterial cell cultures (DH5α) and purified following the PureYield Plasmid MidiPrep System according to the manufacturer’s protocol (Promega, A2492). The concentration and purity of the fresh DNA plasmids were determined using a spectrophotometer (ThermoFisher Scientific, NanoDrop 2000), making sure that their ratio of absorbance fell within the recommended range (A260/A280=1.7–2.0 and A260/A230=2.0 - 2.2) to avoid usage of contaminated samples.

### Cell culture

All cell lines were cultured at 37°C and 5% CO2 (PHCbi CO2 Incubator) within T-75 Nunc EasYFlask cell culture filtered flasks (ThermoFisher Scientific, 156499). Cells were cultured within their respective media supplemented with 10% foetal calf serum (ThermoFisher Scientific, 32561029), 100 units/mL penicillin and 100 µg/mL streptomycin (Gibco). MCF-7 (ECAAC 86012803) and HeLa cell lines were cultured within Gibco Minimum Essential Media (MEM) with GlutaMAX supplement without nucleosides. BT-474 and SKRB3 cell lines were cultured within 25 mM D-glucose and L-Glutamine Dulbecco’s Modified Eagle’s Medium (Gibco). BT-549 and ZR-75 cell lines were cultured within Roswell Park Memorial Institute 1640 medium (Gibco).

Cell density was quantified using an automated cell counter (Bio-Rad TC20TM), while signs of contamination and cellular stress were routinely monitored using an inverted microscope (Motic AE31E Binocular Microscope). Experiments were performed using cells between passages 4 and 25.

### Transient transfection

Cells were plated onto uncoated 13mm glass coverslips and transiently transfected with Halo-*ESR1* constructs (GeneArt ThermoFisher Scientific) using FuGene HD (1.5:1, Promega) in Opti-MEM (Gibco) according to a previously optimised protocol in the Toseland Lab. The transfection mixture was incubated for 20 minutes at room temperature and then added to 24-well plates containing cells at 70-80% confluency. After 5 hours, the transfection mixture was replaced with the corresponding media of each cell line.

Transfected cells were incubated for 24 hours and then labelled with Halo-ligand Tetramethyl Rhodamine (TMR, 1:50000, Promega G8252) and Hoechst 33342 (1:12000, ThermoFisher Scientific, 62249). Cells were fixed with pre-warmed 4% (w/v) paraformaldehyde (PFA) in phosphate-buffered saline (PBS) (ThermoFisher Scientific, J61899AK) for 15 minutes, and the residual PFA was quenched with 50 mM NH4Cl/TBS for 15 minutes. Coverslips were stored within TBS at 4°C until mounting with Mowiol (10% (w/v), 25% (w/v) glycerol, 0.2M Tris-HCl (pH 8.5) supplemented with 2.5% DABCO).

### Drug treatments

To probe the biophysical properties of condensates, transfected and fluorescently labelled cells were treated with 1,6-hexanediol (Sigma-Aldrich, 240117) at final concentration of 2.5%, 5% or 10% to disrupt weak hydrophobic interactions. The 1,6-hexanediol was prepared in imaging media (Gibco Minimum Essential Media without L-glutamine, HEPES or Phenol Red) and filtered before use, with cells imaged immediately using time-lapse live cell microscopy.

To induce chromatin relaxation, cells were treated with the histone deacetylase (HDAC) inhibitor Trichostatin A (TSA, Selleckchem) at 200nM concentration for 24 hours. Upon TSA treatment, cells were fluorescently labelled with Halo-ligand Tetramethyl Rhodamine (TMR, 1:50000, Promega G8252) and Hoechst 33342 (1:12000, ThermoFisher Scientific, 62249). Cells were then either fixed and mounted onto coverslips or used immediately for live-cell optical tweezers experiments.

### Widefield microscopy

Transfection efficiency was assessed using a Zeiss Cell Discoverer 7 widefield microscope with a Plan-Apochromat 20x/0.7 or 5x/0.35 dry objective. Fluorescence imaging was performed using 405nm and 561nm excitation lines for Hoechst 33342 and Halo-ligand TMR, respectively. ZEN Blue 3.1 software was used to acquire images for further analysis on ImageJ (version 1.53q, via Fiji). The percentage of TMR-positive cells relative to total Hoechst-stained nuclei was used to determine transfection efficiency.

### Confocal microscopy

Confocal and Airyscan imaging were performed using a Zeiss LSM 980 microscope with a Plan-Apochromat 63×1.4 NA oil immersion objective (Carl Zeiss, 420782-9900-000). For confocal imaging, fluorescence was excited with 405 nm and 561 nm laser lines, and images were acquired with a pinhole of 1 AU. Airyscan imaging was performed in Super-Resolution mode using the same laser lines, and raw data were processed using the Zeiss Airyscan processing algorithm in ZEN Blue 3.1. All images were analysed in ImageJ (version 1.53q, via Fiji) to quantify the number and area of *ESR1* fusion condensates.

### Live cell imaging

MCF7 cells were cultured in 4-well chambered coverglass dishes with a No.1.5 borosilicate glass bottom (ThermoFisher Scientific 155409, Nunc Lab-Tek II). Upon transfection, cells were incubated for 24 hours and then labelled with Halo-ligand Tetramethyl Rhodamine (TMR, 1:50000, Promega). Cells were maintained in Gibco Minimum Essential Media (MEM) without L-glutamine, HEPES or Phenol Red during imaging (ThermoFisher Scientific, 51200038). Live cell imaging of transfected cells was performed using a ZEISS LSM980 microscope with Airyscan 2 and a Plan Apochromat 63x 1.4 NA oil immersion objective (Carl Zeiss, 420782-9900-000). The microscope was set at 37 °C with 5% CO₂.

### Structured illumination microscopy

Structured illumination microscopy (SIM) was performed using a Zeiss Elyra PS.1 super-resolution imaging system (Carl Zeiss). Images were acquired using a Zeiss Plan-Apochromat 63×/1.4 Oil DIC M27 objective lens. Fluorescence was excited using a 405-nm diode laser for Hoechst staining and a 561-nm diode laser for TMR-labelled samples. For SIM reconstruction, five grid rotations were captured for each optical section. Serial z-stack images were acquired at 0.31um intervals to capture multiple focal planes. Raw SIM data were reconstructed and processed using ZEN Black (Zeiss).

### Condensate analysis

*ESR1* fusion condensates were analysed on ImageJ (version 2.1.0/1.53q) to quantify their area, number, and roundness. Individual nuclei were selected as regions of interest (ROIs) using the Hoechst channel, and condensates were identified by thresholding HaloTag TMR fluorescence. Measurements were performed using the “Analyse Particles” function, with objects outside the ROI or clearly non-condensate particles manually excluded.

Condensate dynamics were evaluated using a simplified Brownian coalescence model. The model assumes a spatially homogeneous population of approximately spherical condensates that undergo instantaneous fusion upon collision, with conservation of total condensed-phase volume and no additional nucleation, dissolution, Ostwald ripening, sedimentation or active transport. Under these assumptions, condensate number follows:

N_t_= N_0_/(1+(t/*tau*))

Where Nt is the number of condensates at time (t), N0 is the initial condensate number and *tau* is the characteristic coalescence timescale. The model was fitted directly to the untransformed condensate-count data by nonlinear least-squares regression.

### Condensate tracking

Time-lapse images of *ESR1* fusion condensates were analysed using the TrackMate plugin in ImageJ (version 1.54r, via Fiji). Images were acquired every 30 seconds for 15-30 minutes and used for condensate tracking. For cells that displayed significant movement, images were pre-aligned using the StackReg plugin to correct for drift. Spot detection was performed using the LoG detector with sub-pixel localisation, and trajectories were generated using the Kalman algorithm. Condensate dynamics, including displacement, speed, and maximum distance travelled, were extracted from TrackMate outputs.

### Fluorescence Recovery After Photobleaching (FRAP)

FRAP experiments were performed on a custom-built microscope with Nikon Eclipse Ti2 stand, equipped with a Cairn multiline laser bank, iLas2 TIRF & scanning FRAP laser illumination module, standard filter sets and Photometrics Prime 95B camera. Cells were imaged using a Nikon 100x Apo TIRF oil objective. Cells were maintained at 37°C in CO2-independent medium without L-Glutamine (Gibco, 18045070) during imaging. Regions of interest (ROIs) within nuclei were photobleached using a 561nm laser at 15% power, with 50 repetition pulses at 25ms per pulse, hitting an effective thickness/depth of approximately 5µm. Fluorescence recovery was recorded at 30-second intervals for 10-20 minutes. All image acquisition and laser targeting were controlled via the MetaMorph software (version 7.10.5.476) and Modular software respectively.

FRAP recovery curves were analysed using the FRAP Analysis plugin in ImageJ (version 1.54r, via Fiji). The photobleached nuclear region was manually selected as the ROI, and the raw fluorescence intensities were normalised using a background reference region and an unbleached control region. Recovery kinetics, including the mobile fraction and half-time (t½), were extracted from the plugin output.

### Optical trapping

Live-cell condensate perturbation experiments were performed using the Lumicks C-Trap system, which combines optical tweezers with confocal fluorescence imaging. Samples were prepared in a flow cell and maintained at room temperature in CO2-independent media without L-Glutamine (Gibco, 18045070) during imaging. Confocal imaging of the condensates was performed using a 561nm laser line and 600/50 nm bandpass filter. A single optical trap was set to move back and forth across the nucleus to probe the mechanical behaviour of condensates. Condensate displacement, speed and coalescence were quantified using the TrackMate plugin in ImageJ (version 1.54r, via Fiji).

### Atomic Force Microscopy (AFM)

Samples were prepared on 35mm WillCo dishes (Ibidi), and cells were maintained at room temperature in CO2-independent media without L-Glutamine (Gibco, 18045070) during imaging. AFM measurements were performed with a Bruker Bioscope Resolve mounted on an inverted microscope (Nikon Eclipse Ti2) connected to an ORCA-Flash4.0LT (Hamamatsu) camera. Pre-calibrated Silicon Tip – Nitride cantilevers (PFQNM-LC-V2 Bruker) were used with a 70nm tip radius. PeakForce Tapping/PeakForce Capture with a maximum indentation force of 0.5 nN was used (43). Indentation curves were fitted within Nanoscope Analysis (Bruker) using a cone-sphere model (44). Three independent regions per sample were analysed.

### Stochastic Optical Reconstruction Microscopy (STORM)

STORM measurements were performed following established protocols (21, 45, 46). Round 25mm No. 1.5 coverslips were treated with etching solution prepared with 5:1:1 distilled water, NH4OH (Sigma-Aldrich, CA, USA) and H2O2 (Thermo Fisher Scientific, MA, USA) at 70°C for 1h. The coverslips were subsequently washed in methanol, followed by distilled water, and then allowed to air dry. The coverslips were exposed to UV for 15 min before plating. Upon transfection, cells were incubated for 24 hours and then labelled with Halo-ligand Tetramethyl Rhodamine (TMR, 1:50000, Promega) and Hoechst 33342 (1:12000, ThermoFisher Scientific, 62249). Cells were fixed using prewarmed 4% (w/v) paraformaldehyde (PFA) in phosphate-buffered saline (PBS) (ThermoFisher Scientific, J61899AK) for 15 minutes, and the residual PFA was quenched with 50 mM NH4Cl/TBS for 15 minutes. Cells were washed with PBS three times before being stored in PBS supplemented with 0.02% w/v sodium azide at 4°C in the dark.

Prior to imaging, the coverslips were washed in PBS twice and placed into an Attofluor cell chamber (Invitrogen, MA, USA). STORM buffer containing 10% w/v glucose, 10 mM NaCl, 50 mM Tris-HCl pH 8.0, with GLOX solution consisting of 5.6% w/v glucose oxidase and 3.4 mg/mL catalase in 50 mM NaCl, 50 mM Tris-HCL pH8.0 and 0.1% w/v β-mercaptoethanol was used for imaging.

STORM images were obtained using a Zeiss Elyra PS.1 system. Sample illumination was achieved using an HR Diode 561 (100mW) laser with 7-12kW/cm2 power density exposed to the sample. The built-in multi-band dichroic mirror MBS 405/561/642 (Carl Zeiss 1784-996) was used to reflect the excitation laser beams onto the sample. Imaging was performed using a 100x NA 1.46 oil immersion objective (Carl Zeiss alpha Plan-Apochromat, 420792-9800-00), and to reduce background fluorescence levels, highly inclined and laminated (HILO) illumination was used. To capture fluorescence signal, a BP420-480/BP 570-640/LP 40 multi-bandpass emission filter (Carl Zeiss 1769-207) was used. The final image acquisition was achieved using an Andor iXon Ultra 897 EMCCD camera with a 30ms exposure for 20000 frames.

Zeiss Zen Black software was used to detect blinking events with a 9-pixel mask and a signal-to-noise ratio of 6. Final molecule positions were determined through fitting a 2D Gaussian function. Model-based cross-correlation drift correction was used on the STORM image. For HaloTag TMR the mean value of localisation procession was 30nm.

### Quantitative analysis of STORM localisation clustering

Single-molecule localisation coordinates generated from STORM reconstructions were exported as localisation tables containing the x- and y-position of each detected event together with localisation precision, photon number and frame information. Subsequent quantitative analyses were performed using a custom Python workflow.

For each nucleus, localisation coordinates were converted from nm to µm and binned onto a two-dimensional grid with a pixel size of 50 nm. Localisation-count maps were smoothed using a Gaussian filter (σ = 1.2 pixels; approximately 60 nm) to generate continuous localisation-density maps. High-density regions were identified independently for each acquisition using a three-component Gaussian mixture model fitted to the log-transformed [log(1+x)] non-zero pixel-density distribution. The component with the highest mean density was classified as the high-density population. Connected high-density regions smaller than 16 pixels (0.04 µm²) were removed, and the remaining connected components were defined as individual ESR1-DAB2 clusters. For each segmented cluster, the area, localisation number, and centroid position were calculated. Cluster area was determined directly from the number of segmented pixels and expressed in µm². The proportion of ESR1-DAB2 localisations associated with clusters was calculated for each nucleus as the sum of localisation events assigned to all segmented clusters divided by the total number of valid nuclear localisations, expressed as a percentage. Cluster number and mean and median cluster area were calculated independently for each nucleus.

### Radial localisation-density analysis of ESR1-DAB2 clusters

To determine whether ESR1-DAB2 assemblies displayed internal nanoscale organisation, radial localisation-density profiles were calculated from segmented clusters. To minimise effects arising from irregular, overlapping or poorly sampled structures, clusters were included in the radial analysis when they had an area ≥0.05 µm², contained ≥150 localisation events, had an eccentricity ≤0.85 and solidity ≥0.80, and were spatially isolated from neighbouring clusters. Spatial isolation was defined using the ratio between nearest-neighbour centroid distance and cluster equivalent radius, with clusters retained when this ratio was ≥3.0.

For each retained cluster, an equivalent circular radius (REQ) was calculated from its segmented area:

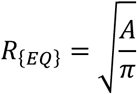

where (*A*) represents cluster area. The geometric centroid of the segmented cluster was used as the origin, and the radial distance of each localisation from the centroid was expressed relative to REQ. Localisations were assigned to concentric radial intervals of 0.25 REQ, extending from the cluster centre to 1.75 REQ. Localisation density within each radial interval was calculated by dividing the number of localisation events by the corresponding annular area and expressed as localisations µm⁻². This area correction accounts for the increasing area of successive radial annuli. Radial positions between 0 and 1.0 (REQ) were considered to represent the internal cluster region, whereas positions beyond 1.0 (REQ) sampled the surrounding environment. Local nucleoplasmic background density was estimated from the combined 1.25–1.75 (REQ) region. Localisation enrichment was calculated as the ratio between localisation density within each radial interval and the corresponding local background density. To allow comparison between clusters with different absolute localisation densities, profiles were additionally normalised to the maximum localisation density measured within the four internal radial intervals. Differences in localisation density across the four internal radial intervals (0–0.25, 0.25–0.50, 0.50–0.75 and 0.75–1.00 REQ) were assessed using a Friedman test. Consecutive radial intervals were subsequently compared using paired one-sided Wilcoxon signed-rank tests with Holm correction for multiple comparisons.

### Fluorescence-lifetime imaging microscopy (FLIM)

MCF-7 cells were cultured in 4-well chambered coverglass dishes with a No. 1.5 borosilicate glass bottom (ThermoFisher Scientific, Nunc Lab-Tek II). Following transfection, cells were labelled with Halo-ligand Tetramethyl Rhodamine (TMR, 1:50000; Promega). Cells were maintained at 37°C in CO2-independent medium without L-Glutamine (Gibco) during imaging.

FLIM measurements were performed on a ZEISS LSM980 laser-scanning confocal microscope equipped with an Airyscan 2 module and a Plan-Apochromat 63x/1.4 NA oil-immersion objective, controlled via ZEN blue 3.7 software. Time-Correlated Single Photon Counting (TCSPC) data acquisition was managed using an integrated PicoQuant MultiHarp detection system operated in T3 mode.

Samples were excited with a 561 nm laser at 100% power and a repetition rate of 40 MHz. Images were acquired in unidirectional scan mode using a 512 x 512 pixel frame size and a scan speed setting of 4. The confocal pinhole was set to 1 Airy unit, and fluorescence was detected using a GaAsP photomultiplier tube. A maximum acquisition time of 5 min per field of view was used to standardise photon collection.

FLIM datasets and spatial lifetime maps were processed and analysed using SymPhoTime 64 (PicoQuant). Fluorescence lifetime decays over a temporal range of 0.08–24.91 ns were evaluated using Fast FLIM with a mono-exponential reconvolution model (n = 1) and a calculated instrument response function (IRF). Data were processed using a spatial binning factor of 2. An initial fit was performed to determine baseline parameters, after which the background decay, IRF shift and background IRF were set to zero. During the final fit, the lifetime (τ1) and amplitude (A1) parameters were allowed to vary to extract quantitative fluorescence lifetimes. The resulting fluorescence lifetime maps were used to define regions of interest (ROIs) corresponding to the interior of *ESR1* fusion condensates and the surrounding diffuse nucleoplasm and fitted τ1 values were extracted for comparison.

### Slimfield microscopy

Slimfield microscopy (25) was performed using a bespoke single-molecule super-resolved microscope with millisecond temporal resolution when imaging fluorescently tagged proteins. The microscope is built around an openFrame microscope body (Cairns Research), a Nikon CFI Apo TIRF 100X NA1.49 Oil objective and nano-LP (Mad City Labs) 3-axis nano-positioning stage. A 50 mW 561 nm laser (Coherent OBIS LS 561 nm 50 mW Laser System) is relayed through a 1:1 Keplerian telescope and focussed onto the back focal plane of the objective lens (Nikon CFI Apo TIRF 100X NA1.49 Oil), generating an epifluorescence excitation of ∼12 µm at FWHM at the sample. Emitted fluorescence is then transmitted to a TeleDyne Photometrics Prime BSI sCMOS camera via an Optosplit (Cairns Research) with a magnification of 1.28X resulting in an imaging pixel sixe of 51 nm. Back reflected laser light was removed from the signal with a 75 nm band pass filter centred on 610 nm. Stepwise photobleaching is used to identify single fluorescent proteins (47), and this information can then be used to assign high precision estimates of stoichiometry to molecular complexes and assemblies such as condensates (47, 48).

For imaging condensate size, the laser power was set to 12 mW and attenuated to 1.2 mW using an ND1 filter, giving a power density at the sample of ∼830 W/cm^2^. The laser was controlled via digital modulation by the sCMOS camera, preventing any unnecessary photobleaching, with a frame-to-frame time of 5 ms.

Subsequent imaging of single *ESR1* fusion proteins for diffusivity measurements was performed with the 561 nm laser set to 12 mW, giving a power density at the sample of ∼8300 W/cm^2^, with the laser digitally modulated by the sCMOS camera. Images were acquired with a frame-to-frame time of 5 ms.

### Single-particle tracking analysis

Single-particle tracking analysis was performed on fluorescence microscopy image sequences using the Python-based software package PySTACHIO (49). Fluorescent spots corresponding to individual labelled molecules were detected in each frame and linked across successive frames to generate particle trajectories. Tracking parameters included a maximum displacement of 5 pixels between frames, a minimum trajectory length of 2 frames, Gaussian image filtering, a frame interval of 5ms, and a pixel size of 0.051μm. The resulting trajectories were subjected to post-processing analysis within PySTACHIO to determine diffusion coefficients and stoichiometry.

### IDR prediction tools

Protein sequence analysis was performed to assess amino acid composition, intrinsic disorder, and phase separation propensity of *ESR1* fusion proteins. The ExPASy ProtParam tool (50) was used to determine the amino acid composition and grand average of hydropathy (GRAVY) scores based on the Kyte-Doolittle hydropathy scale. Protein intrinsic disorder was evaluated using the Predictor of Natural Disordered Regions (PONDR) web server (51), using the PONDR VSL2 predictor algorithm to characterise both short and long disordered segments. To further evaluate the phase separation potential, sequences were analysed using the ParSe v2 algorithm (52), which classifies residues into phase-separating, disordered non-phase-separating, and structured regions based on sequence composition and patterning.

### Statistics

All quantitative data are presented as mean ± standard deviation (SD) from at least three independent biological replicates, unless otherwise stated. Statistical significance between two groups was evaluated using the non-parametric Mann-Whitney *U* test. For comparisons among three or more groups, the Kruskal-Wallis test followed by Dunn’s post-hoc test for multiple comparisons was performed. A *p*-value < 0.05 was considered statistically significant (*ns, not significant; *p < 0.05, **p < 0.01, ***p < 0.001, ****p < 0.0001*). All statistical analyses and curve fitting were performed using GraphPad Prism (Version = 11.0.1).

### Graphics

Unless stated, data fitting and plotting were performed using GraphPad Prism (Version = 11.0.1). Diagrams were generated using the BioRender software.

### Data Availability

The data supporting the findings of this study are available from the corresponding author on request.

## RESULTS

### *ESR1* fusion proteins exhibit increased intrinsic disorder following structural rearrangement

The generation of *ESR1* fusions alters the composition and predicted structural properties of the resulting chimeric proteins. To characterise these changes, we analysed a panel of recurrent *ESR1* fusion variants and compared their amino acid composition, hydropathy and predicted intrinsic disorder with wild-type *ESR1* (*WT-ESR1*).

Several *ESR1* fusion proteins were enriched in residues associated with intrinsic disorder and low-complexity regions, including proline, serine, glycine and alanine (Figure 1A, Supplementary table 1). Conversely, residues typically associated with structured globular domains, including cysteine and tryptophan, were comparatively underrepresented in most fusions (Supplementary figure 1A). *ESR1-GYG1* and *ESR1-MTHFD1L* were notable exceptions, consistent with these partner proteins being more structured.

**Figure 1:**
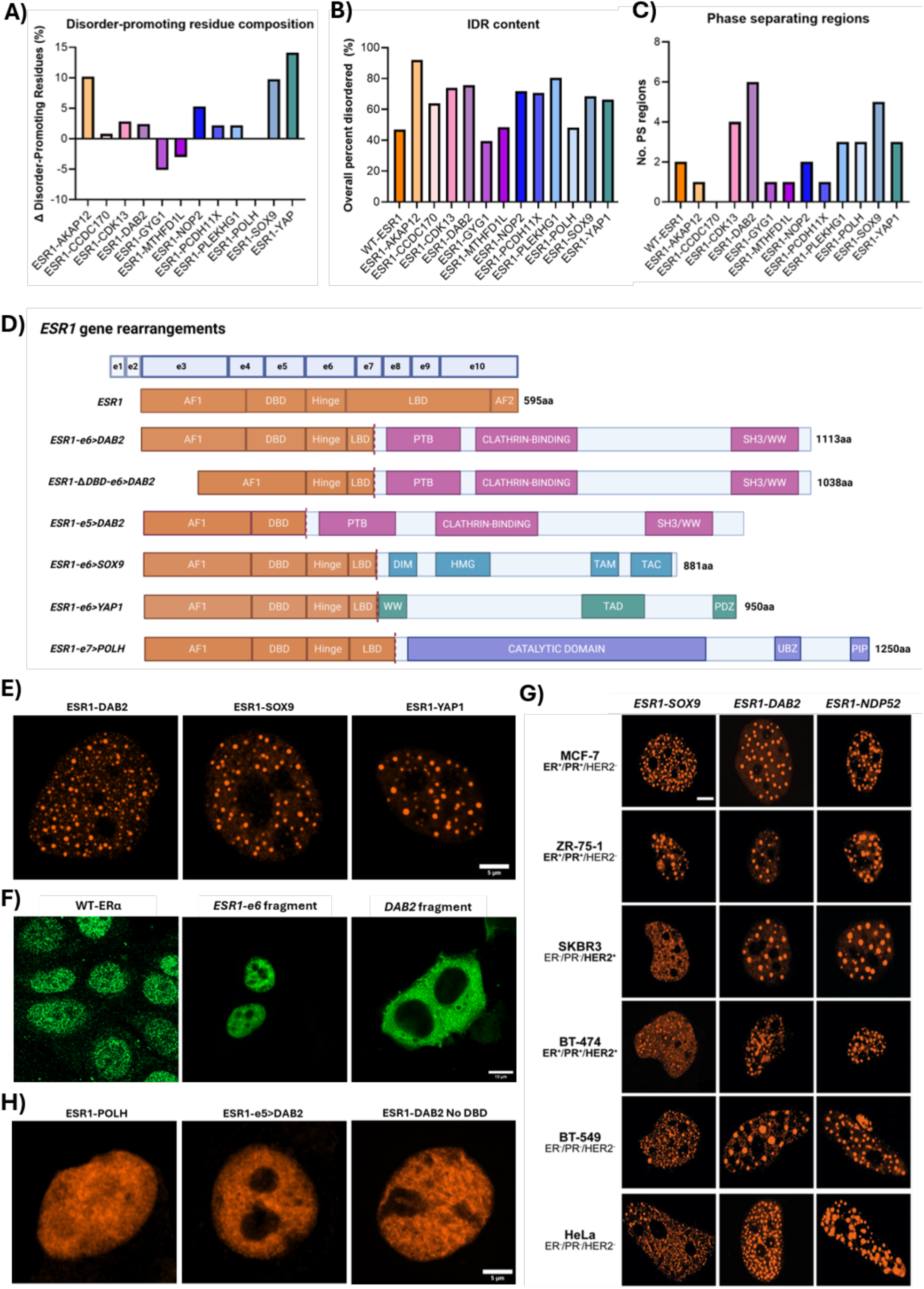
Sequence and structural determinants of nuclear foci formation by ESR1 fusion proteins. (A-C) Sequence analysis of ESR1 fusions compared to wild type ESR1 (WT-ESR1). **(A)** Difference (Δ) in percentage of disorder-promoting residue composition relative to wild-type ESR1, including alanine (A), glutamic acid (E), glycine (G), lysine (K), proline (P), glutamine (Q), arginine (R), and serine (S) which represent flexible, polar or charged amino acids. **(B)** Total predicted intrinsically disordered region (IDR) content (%) determined using the PONDR VSL2 prediction tool. **(C)** Number of phase-separating regions for each ESR1 fusion variant compared to WT-ESR1, calculated using the ParseV2 prediction tool. **(D)** Schematic structural architecture of the ESR1 gene and its fusion variants. ESR1-DAB2, ESR1-SOX9, and ESR1-YAP1 integrate at exon 6 of ESR1, whereas ESR1-POLH fuses at exon 7, both resulting in the truncation and loss of the C-terminal ligand-binding domain (LBD). The engineered ESR1-e5>DAB2 variant was designed so that DAB2 is fused to exon 5 instead of exon 6 of ESR1. The ESR1-DAB2 No DBD variant represents a targeted deletion construct designed to eliminate the functional DNA-binding domain (DBD) while preserving the remainder of the fusion sequence. **(E)** ESR1-e6> fusions recurrently form nuclear foci. Airyscan images of MCF7 cells transfected with various ESR1 fusion variants (ESR1-DAB2, ESR1-SOX9, ESR1-YAP1) and labelled with Halo-ligand TMR. Scale bar = 5μm across all panels. **(F)** Confocal images of MCF-7 cells immunostained for ERα or transiently expressing individual truncated fragments (ESR1-e6> fragment or >DAB2 fragment) and fluorescently labelled with Halo-ligand Oregon Green. Scale bar = 10μm across all panels. **(G)** Confocal images of breast cancer lines (MCF-7, ZR-75-1, SKBR3, BT-474, BT-549) and the cervical cancer line HeLa, transiently transfected with the Halo-tagged fusions ESR1-DAB2, ESR1-SOX9 or the pseudo-fusion control ESR1-NDP52. Fusions were fluorescently labelled using the Halo-ligand TMR. Scale bar = 5μm across all panels. **(H)** Airyscan images of MCF7 cells transfected with ESR1-POLH, ESR1-e5>DAB2 or ESR1-DAB2 No DBD and labelled with Halo-ligand TMR. Scale bar = 5μm across all panels. **Abbreviations:** aa, amino acids; AF1, activation function 1; AF2, activation function 2; DBD, DNA-binding domain; DIM, dimerization domain; e (e1–e10), exon; HMG, high-mobility group; Hinge, hinge domain; LBD, ligand-binding domain; PDZ, PDZ domain-binding motif; PIP, PCNA-interacting peptide motif; PTB, phosphotyrosine-binding domain; SH3/WW, Src homology 3 and WW domain-binding region; TAD, transactivation domain; TAM, transactivation motif; TAC, transactivation C-terminal domain; UBZ, ubiquitin-binding zinc finger domain; WW, tryptophan-tryptophan domain.

Consistent with these compositional changes, PONDR predictions showed increased intrinsic disorder across most *ESR1* fusions, except for *ESR1-GYG1* and *ESR1-MTHFD1L* (Figure 1B). These findings suggest that the extent of disorder is largely dictated by the fusion partner, particularly in variants incorporating highly disordered proteins (Supplementary figure 1B).

Hydropathy analysis further revealed that most *ESR1* fusions had more negative GRAVY scores than *WT-ESR1*, indicating a shift towards hydrophilicity (Supplementary figure 1C). This was accompanied by depletion of hydrophobic and aromatic residues and, in several cases, enrichment of charged residues (Supplementary table 1), consistent with sequence features typical of intrinsically disordered regions.

We next assessed whether increased disorder was associated with phase-separation propensity by analysing the fusion sequences using ParSe_v2. While *WT-ESR1* contained two predicted phase-separating regions, several fusions, including *ESR1-DAB2*, *ESR1-SOX9* and *ESR1-YAP1* showed expanded phase-separating architecture (Figure 1C). In contrast, *ESR1-CCDC170* showed little predicted phase-separating potential, further supporting the idea that distinct fusion partners impose different biophysical properties on the *ESR1* fusion protein.

Collectively, these analyses indicate that many *ESR1* fusions are structurally distinct from *WT-ESR1*, characterised by increased intrinsic disorder, hydrophilicity and predicted phase-separation propensity. By combining the retained AF-1/DBD region of ERα with partner-derived disordered domains, these chimaeras may gain expanded interaction interfaces and enhanced multivalency. These properties provide a structural basis for higher-order nuclear assembly and suggest that *ESR1* fusions may couple chromatin targeting with condensate-forming potential.

### *ESR1* fusions recurrently form nuclear protein clusters in a variety of breast cancer cell lines

To investigate how these structural changes and increased disorder may impact protein distribution, we next assessed the intracellular organisation of *ESR1* fusion proteins. The breast cancer cell line MCF-7 was transiently transfected with constructs encoding the *ESR1-e6>DAB2*, *ESR1-e6>SOX9* and *ESR1-e6>YAP1* fusions (Figure 1D). These specific variants were selected because they are recurrent and exhibit some of the most pronounced structural and compositional changes in the dataset, driven by their high intrinsic disorder and phase-separating potential. Furthermore, the fusion partners are functionally distinct, whereby *SOX9* and *YAP1* act as transcription factors, while *DAB2* is an adaptor protein involved in endocytosis. Therefore, this allows us to evaluate whether their shared structural characteristics drive similar intracellular behaviour despite their different native roles.

Upon transfection, all three fusions formed distinct, well-defined nuclear foci that were highly variable in size and number (Figure 1E). In contrast, wild-type ERα (WT-ERα) or the individual fusion fragments alone did not produce the same organisation and distribution (Figure 1F), indicating that cluster formation is a fusion-specific property arising from the rearrangement of *ESR1* with another gene. While WT-ERα and the *ESR1-e6>* fragment can also localise in small punctate structures (18), the *ESR1* fusion clusters were markedly different, appearing as larger, more prominent assemblies, highlighting a more exaggerated phenotypic shift in protein distribution.

To confirm that this clustering is not a cell line-specific event, a panel of breast cancer cell lines was transiently transfected with *ESR1-e6>DAB2* and *ESR1-e6>SOX9*. Both fusions consistently concentrated into nuclear clusters with comparable morphology and distribution across all cell lines tested (Figure 1G). This phenotype was further observed in the cervical cancer cell line HeLa, demonstrating that the propensity to concentrate into these nuclear assemblies is an intrinsic property of *ESR1* fusion proteins that occurs independently of cell type or hormone receptor status. Based on this consistency, all subsequent experiments were performed in the ER-positive MCF-7 breast cancer cell line, which serves as a well-established and disease-relevant model.

### Exon 6 rearrangement facilitates higher-order assembly of *ESR1* fusion proteins

The majority of *ESR1* fusion breakpoints occur after exon 6, which eliminates most of the ligand-binding domain (LBD) while retaining the DNA-binding domain (DBD) (Figure 1D) (11, 19, 20). To determine whether the formation of nuclear foci are driven by the *ESR1-e6>* configuration, we engineered a *de novo* fusion, *ESR1-e6>NDP52*. This variant is not found in patients but contains the *NDP52/CALCOCO2* gene, which is inherently highly disordered (21). Notably, this artificial fusion also formed robust nuclear assemblies across all cell lines tested (Figure 1H), indicating that higher-order assembly is a more generalised property of the *ESR1-e6>* fusion architecture, particularly when incorporating intrinsically disordered C-terminal partners.

Alternative *ESR1* fusion breakpoints involving exons 4, 5 and 7 have also been observed in patient samples, although they are rarer (9, 11). To assess whether these alternative structural rearrangements similarly confer the capacity to cluster, we evaluated a series of additional *ESR1* fusion variants in MCF-7 cells. The fusions *ESR1-e5>DAB2* and *ESR1-e7>POLH* generated at exon 5 or exon 7, respectively, did not form nuclear foci and instead exhibited a diffuse nucleoplasmic distribution (Figure 1H). These findings highlight the importance of the exon 6 breakpoint in facilitating higher-order assembly and that truncation of *ESR1* alone or disruption at adjacent exons is insufficient to induce the formation of these distinct nuclear foci.

### The *ESR1* DNA-binding domain is a critical determinant of higher-order nuclear assembly

The change in the spatial organisation of *ESR1* fusion proteins in the presence of highly disordered domains prompted us to examine which other structural elements may be contributing to this novel phenotype. Depending on the partner gene, *ESR1* fusions acquire or lose various functional domains because of the rearrangement; however, the *ESR1* DBD is conserved across most fusion classes (Figure 1D) (9, 11, 19). To assess whether the DBD influences the nuclear distribution of *ESR1* fusion proteins, we deleted this domain from the *ESR1-e6>DAB2* fusion. Notably, the loss of the DBD abolished cluster formation (Figure 1H), resulting in a diffuse nuclear distribution instead. This suggests that interaction with chromatin may be a necessary scaffold to facilitate higher-order assembly of *ESR1* fusion proteins.

### *ESR1* fusion proteins form spherical condensates consistent with liquid-liquid phase separation

Transcription factor fusion oncoproteins can organise into higher-order nuclear assemblies through mechanisms associated with liquid-liquid phase separation (LLPS) (14, 16, 22). To determine whether *ESR1* fusion proteins form structures with condensate-like properties, we performed a quantitative analysis of foci morphology in MCF-7 cells expressing *ESR1-e6>DAB2*, *ESR1-e6>SOX9* or *ESR1-e6>YAP1*.

Across all three fusions, nuclear assemblies displayed high morphological diversity (Figure 2A), ranging from 1 to 148 condensates per cell and from 0.04 μm² to 5.31 μm² in area. With respect to the latter, the lower limit is constrained by the optical resolution of the microscope. *ESR1-DAB2* formed the highest number of foci per cell (mean=36), followed by *ESR1-YAP1* (mean=22) and *ESR1-SOX9* (mean=19) (p <0.0001 for both comparisons; Figure 2B). In terms of size, *ESR1-SOX9* (mean = 0.68μm²) and *ESR1-DAB2* (mean = 0.48 μm²) formed the largest assemblies and did not differ significantly from one another (p = 0.59) (Figure 2C). Conversely, *ESR1-YAP1* assemblies were significantly smaller on average (mean = 0.45 μm²) compared to both *ESR1-SOX9* (p < 0.0001) and *ESR1-DAB2* (p = 0.0002).

**Figure 2:**
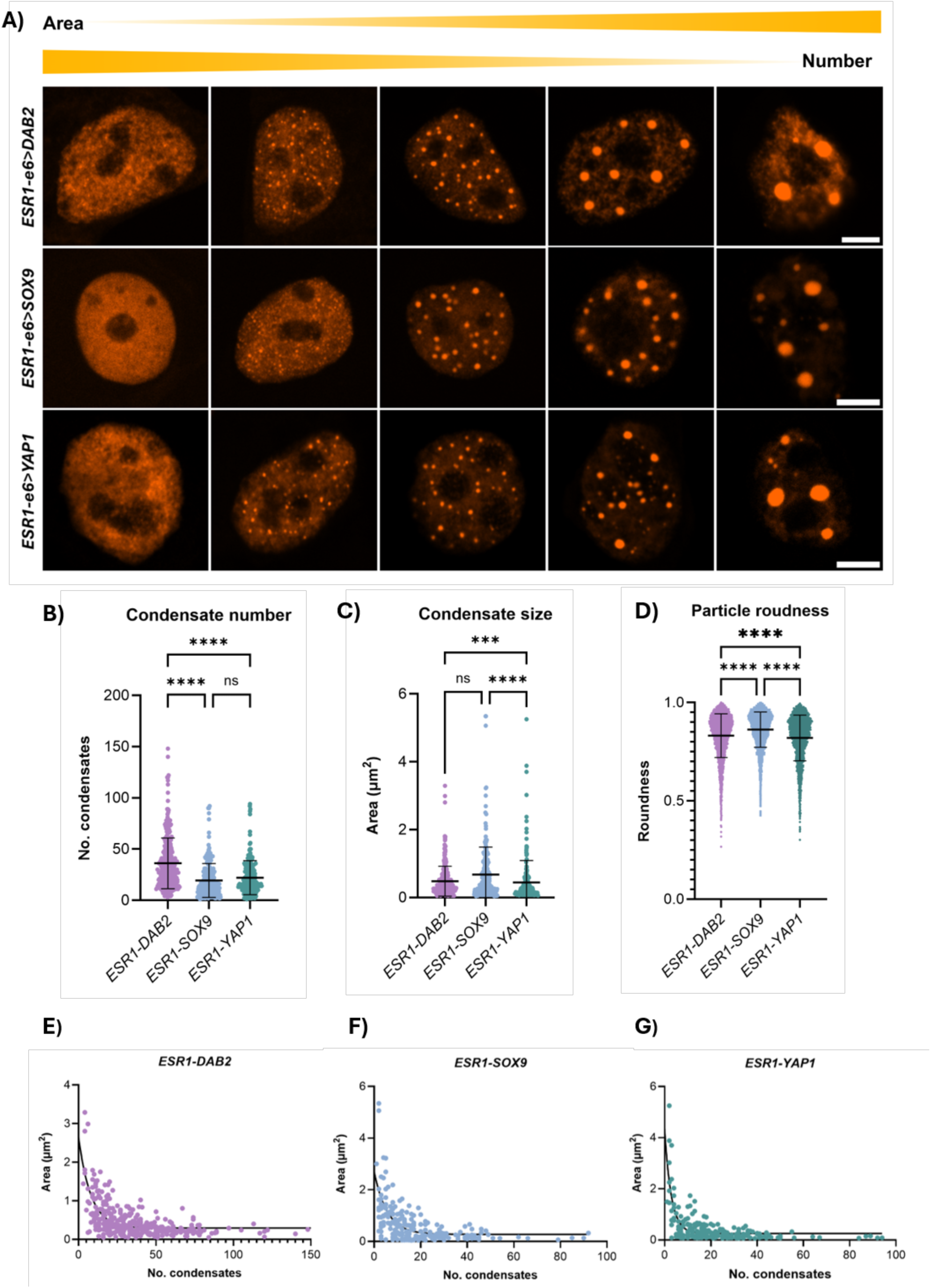
Morphological characterisation reveals progressive coalescence of ESR1 fusion assemblies. **(A)** Morphological variability and maturation stages of ESR1 fusion foci. Representative Airyscan images of MCF-7 cells transfected with Halo-tagged ESR1-DAB2, ESR1-SOX9, or ESR1-YAP1 and labelled with Halo-ligand TMR. Individual panels capture distinct stages of higher-order assembly, transitioning from a diffuse distribution to small, medium, and ultimately large assemblies. An inverse relationship is observed where cells containing a high number of puncta exhibit smaller individual foci, compared to cells containing fewer, larger foci. Scale bars = 5µm across all panels. **(B-D)** Morphological characterisation of ESR1 fusion assemblies. **(B)** Condensate Number: this ranged from 1 to 148 per cell, with a mean number (± SD) of 36.0 ± 24.7 for ESR1-DAB2, 19.0 ± 16.6 for ESR1-SOX9, and 22.0 ± 16.6 for ESR1-YAP1 (Kruskal-Wallis test, p<0.0001). **(C)** Condensate Area: this ranged from 0.05µm^2^ to 5.35µm^2^ across all variants. The mean area (±SD) was 0.48 ± 0.44µm^2^ for ESR1-DAB2, 0.68 ± 0.81µm^2^ for ESR1-SOX9, and 0.45 ± 0.65µm^2^ for ESR1-YAP1 (Kruskal-Wallis test, p < 0.0001). **(D)** Foci Roundness: Roundness values ranged from 0.27 to 1.00 (1.00 = representing perfect sphericity), with mean values (±SD) of 0.83 ± 0.11 for ESR1-DAB2, 0.86 ± 0.09 for ESR1-SOX9, and 0.82 ± 0.12 for ESR1-YAP1 (Kruskal-Wallis test, p < 0.0001). Graphs display individual cellular data points with lines representing the mean ± SD. Statistical significance evaluated using the Kruskal-Wallis test, followed by Dunn’s post-hoc multiple comparisons test between individual groups. **(E-G)** Correlation between the area and number of ESR1 fusion assemblies. Correlation plots are shown for **(E)** ESR1-DAB2 (R^2^ = 0.4408), **(F)** ESR1-SOX9 (R^2^ = 0.3953), and **(G)** ESR1-YAP1 (R^2^ = 0.5374), modelled using a one-phase exponential decay curve. Data were collected from three biological replicates (n = 318 cells for ESR1-DAB2, n = 241 cells for ESR1-SOX9, and n = 215 cells for ESR1-YAP1).

Based on these distributions, we identified three main populations of cells: (1) those with few but large assemblies (n < 20, area > 2.5 μm²), (2) those with many small puncta (n > 100, area < 0.5 μm²), and (3) an intermediate group. Further analysis revealed an inverse relationship between condensate number and size: cells with fewer clusters tended to exhibit larger foci area, in comparison to cells with many puncta that were smaller in size (Figure 2A, 2E-G). This trend suggests a defined progression in which *ESR1* fusion proteins begin in a diffused state, transitioning into small puncta that progressively grow and merge into larger assemblies. Such maturation and growth patterns are highly consistent with the LLPS model (23).

We next assessed foci shape, as rounded morphology is a common feature of liquid-like condensates (24). Roundness analysis confirmed that the assemblies were generally round, with high mean roundness values of 0.83 for *ESR1-DAB2*, 0.86 for *ESR1-SOX9*, and 0.82 for *ESR1-YAP1* (roundness of 1.0= perfect sphere) (Figure 2D). While these values indicate a high degree of roundness, it is worth noting that measurements for the smallest puncta may be slightly underestimated due to limitations in particle detection.

Collectively, these quantitative morphological features strongly support the classification of *ESR1* fusion clusters as phase-separated condensates. Given these similarities across the different variants, we selected *ESR1-DAB2* as our representative model for all subsequent characterisation experiments.

### *ESR1* fusion proteins organise into high-density macromolecular condensates

To examine the organisation of ESR1 fusion proteins at the nanoscale, we performed stochastic optical reconstruction microscopy (STORM) of Halo-TMR-labelled ESR1-DAB2 in MCF-7 cells (Figure 3A). STORM provides single-molecule localisation information, enabling the molecular organisation underlying the nuclear foci to be resolved.

**Figure 3:**
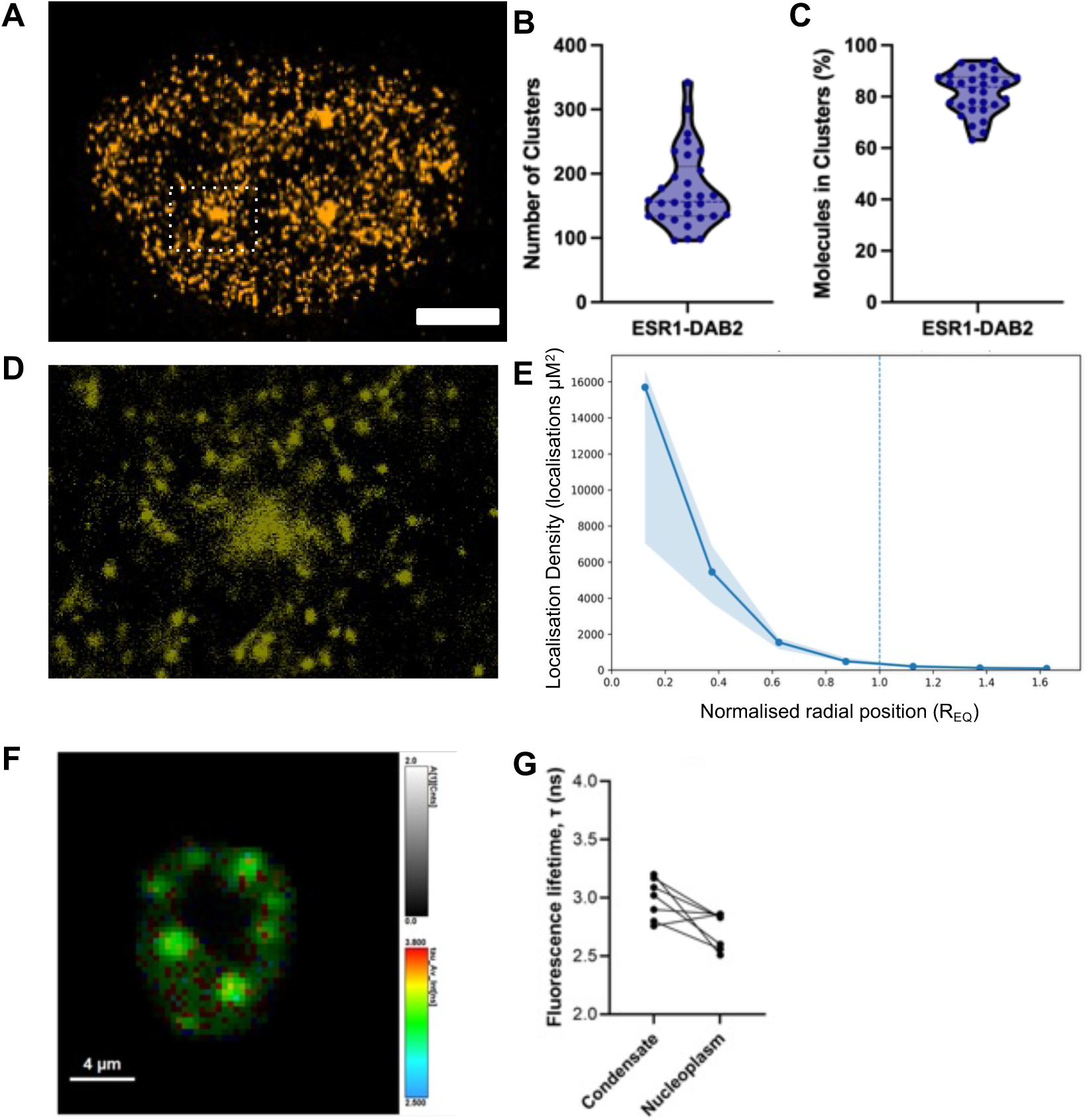
Nanoscale distribution and density of ESR1 fusion proteins. (**A)** Representative STORM render image showing the spatial distribution of HaloTMR-ESR1 fusion proteins in MCF-7 cells. The full-scale overview is rendered by localisation precision to illustrate overall protein organisation. Scale bar = 2μm. (**B)** Quantification of the number of clusters ESR1-DAB2 in a field of view. (**C)** Quantification of the number of ESR1-DAB2 molecules within a cluster. **(D)** Magnified view of the cropped region highlights the localised molecules rendered by plotting individual centroids for each localisation, revealing the exact nanoscale density and cluster organisation of the fusion proteins. **(E)** Radial localisation-density profiles were generated from the STORM acquisitions. For each condensate, localisation density was calculated within concentric radial intervals of 0.25 equivalent radii (REQ), where (REQ) represents the radius of a circle with an area equivalent to that of the segmented condensate. Radial position was normalised such that (r/REQ=0) represents the condensate centre and (r/REQ=1) the condensate boundary (vertical dashed line). The solid line represents the median localisation, with shading indicating the interquartile range (25th–75th percentile). Localisation density differed significantly across the four internal radial intervals (Friedman test, χ² = 33.0, p = 3.22 × 10⁻⁷). Post-hoc paired Wilcoxon signed-rank tests identified significant decreases between each successive internal radial interval after Holm correction for multiple comparisons (adjusted (p=0.00147) for each comparison. (n=30 from three biological repeats). **(F)** Representative FLIM lifetime map of a TMR-labelled ESR1 fusion-expressing MCF-7 cell, with fluorescence lifetime displayed using a fixed pseudocolour scale of 2.5–3.8 ns. Scale bar = 4μm. **(G)** Mean fluorescence lifetime (τ) of condensate and corresponding nucleoplasmic ROIs for individual cells, with paired measurements connected by lines. For each cell, fluorescence lifetime was calculated from three condensate ROIs and three corresponding nucleoplasmic ROIs. Data were retrieved from two biological repeats (n = 7 cells).

Quantitative analysis of STORM localisation distributions revealed that the majority of detected ESR1-DAB2 was partitioned into high-density nuclear assemblies. Across the nuclei, a mean of 83.9% of ESR1-DAB2 localisations were assigned to segmented clusters (Figure 3C), with individual nuclei ranging from 63.1% to 94.0%. These measurements indicate that ESR1-DAB2 is strongly partitioned from the diffuse nucleoplasmic pool into spatially concentrated macromolecular assemblies.

The clusters could be observed as dense assembles within the centre which because more diffuse at the edge. To quantitatively define the nanoscale organisation of ESR1-DAB2 assemblies, we performed radial localisation-density analysis. High-density domains were identified independently within each acquisition using density-based segmentation followed by conservative morphological and spatial filtering. This identifies condensates for analysis. ESR1-DAB2 localisation density was maximal within the central region of the condensates and progressively decreased towards their periphery. Beyond the boundary, localisation density continued to decline and approached the surrounding nucleoplasmic background (Figure 3E). Localisation density differed significantly across the four internal radial intervals, with significant decreases between each successive radial interval. Together, these measurements demonstrate that ESR1-DAB2 condensates possess a reproducible nanoscale architecture characterised by a highly enriched central region and progressively decreasing localisation density towards the condensate periphery.

Importantly, the data therefore provide direct evidence that the nuclear foci observed by confocal microscopy correspond to spatially confined, high-density assemblies of ESR1 fusion molecules. The nanoscale separation between these enriched domains and the surrounding lower-density nucleoplasmic population is consistent with partitioning of ESR1-DAB2 into a condensed molecular state.

Having established that ESR1 fusion foci represent nanoscale regions of elevated molecular density, we next asked whether condensation was accompanied by a change in the local physicochemical environment. We therefore used fluorescence lifetime imaging microscopy (FLIM) to compare Halo-TMR fluorescence within ESR1-DAB2 condensates with the surrounding nucleoplasm. (Figure 3F-G). Analysis of the fluorescence lifetime decay curves revealed that the average fluorescence lifetime within *ESR1* fusion condensates was prolonged by approximately 0.31 ns compared to the diffuse nucleoplasmic phase (Figure 3G; mean τ ± SD = from 2.7 ± 0.22 ns in the nucleoplasm to 3.02 ± 0.2 ns in condensates). This substantial shift in fluorescence lifetime points to a change in molecular crowding, viscosity and/or polarity, confirming that the condensates provide a distinct physicochemical environment. Together, these findings provide evidence that *ESR1* fusion proteins establish specialised, chemically distinct microenvironments through liquid-liquid phase separation.

### *ESR1* fusion condensates form both dynamic and stable membraneless compartments with size-dependent mobility

To further investigate the growth dynamics of *ESR1* fusion condensates, we performed time-lapse live-cell imaging of transfected MCF7 cells (Figure 4A). Distinct fusion events were observed in which smaller condensates coalesced into larger assemblies. While fusion events were relatively infrequent, the coalescence process was rapid once contact was made, with the independent structures merging into a single round body in a manner characteristic of classic liquid droplets (Figure 4B).

**Figure 4:**
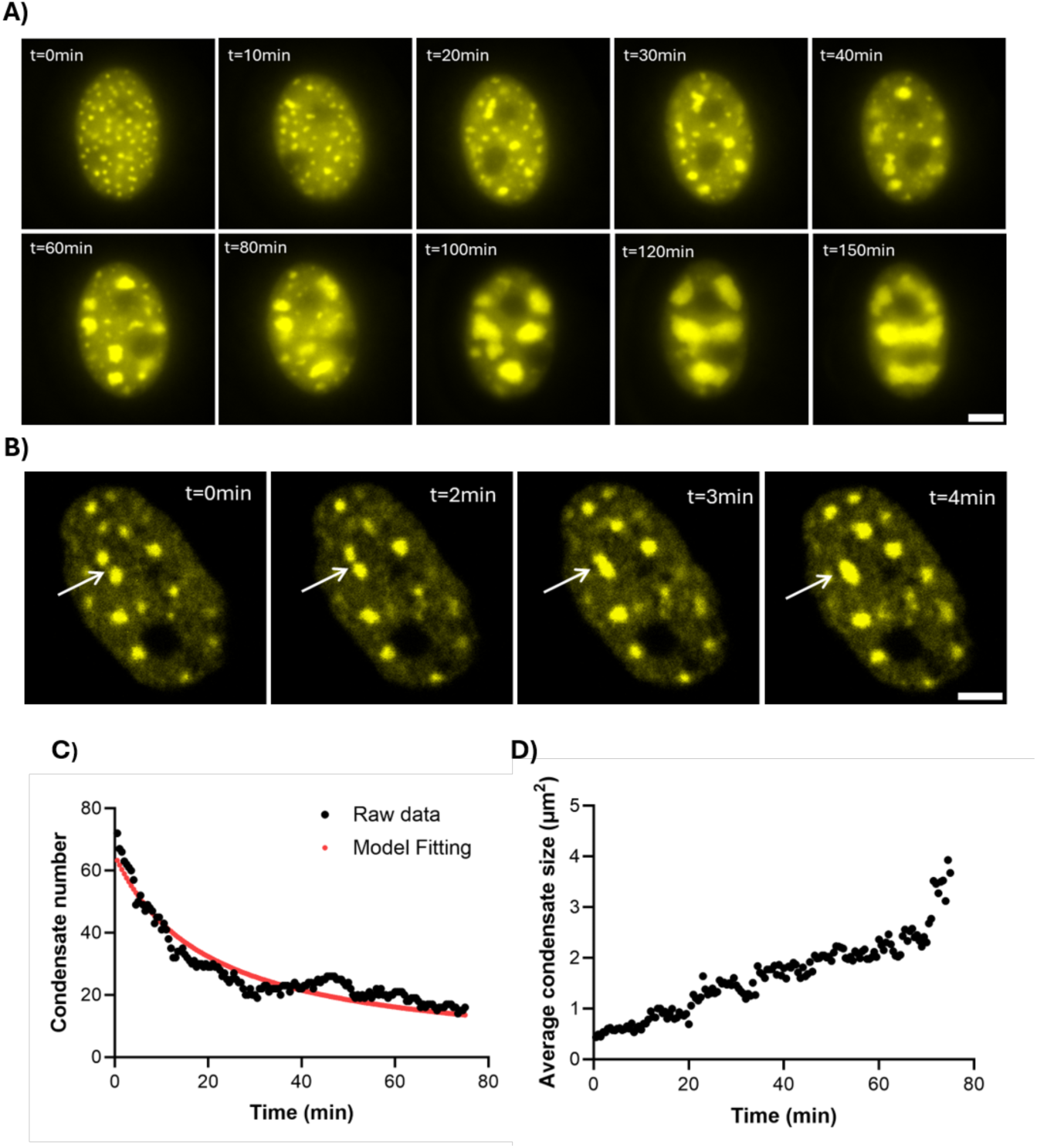
Tracking of ESR1-DAB2 condensate growth, coalescence and mobility. **(A)** Time-lapse live-cell confocal imaging capturing the growth dynamics of ESR1-DAB2 fusion condensates over a 150-minute trajectory. Images show the progressive reduction in condensate number and simultaneous area increase of assemblies over time (t). Scale bar = 5µm across all panels. **(B)** Live-cell time-lapse tracking capturing distinct droplet coalescence of ESR1-DAB2 condensates over time (t). Images highlight the liquid-like properties of the assemblies as independent structures make physical contact and merge into a single, unified condensate. Scale bar = 5µm across all panels. **(C-D)** Quantitative analysis of condensate dynamics over time within a single representative cell showing the change in **(C)** condensate area (µm^2^) and **(D)** condensate number over a 75-minute imaging window.

To quantify these assembly mechanics, we monitored the average size and total number of condensates over a 75-minute trajectory at 30-second intervals (Figure 4C-D). The total number of condensates steadily decreased over time in the first thirty minutes, after which the condensate count plateaued (Figure 4C). Changes in condensate number were analysed using a simplified Brownian coalescence model, which assumes that spherical condensates diffuse, collide and fuse while conserving total condensed-phase volume. Concurrently, the condensate area displayed a steady increase over time (Figure 4D), expanding the average condensate area from around 0.5µm^2^ initially to 4µm^2^ by the end of the trajectory. Taken together, the simultaneous reduction in droplet numbers paired with this accelerating growth profile indicates that condensate enlargement is driven by a cooperative mechanism of distinct droplet coalescence and continuous molecular recruitment.

Condensate dynamics were further quantified using TrackMate (Figure 5A-C). This revealed a correlation between condensate area and mobility: smaller puncta (<0.5µm*^2^*) travelled greater maximum distances (mean ± SD = 0.65 ± 0.29µm) within the nucleus than larger assemblies (>1.5µm*^2^*), which were more static/stationary with restricted and confined movement (mean ± SD =0.36 ± 0.09µm; p = 0.0062). This size-dependent shift from dynamic to stable behaviour may reflect a progressive transition toward a more gel-like or solid-like state over time, which is consistent with the condensate maturation often observed in LLPS.

**Figure 5:**
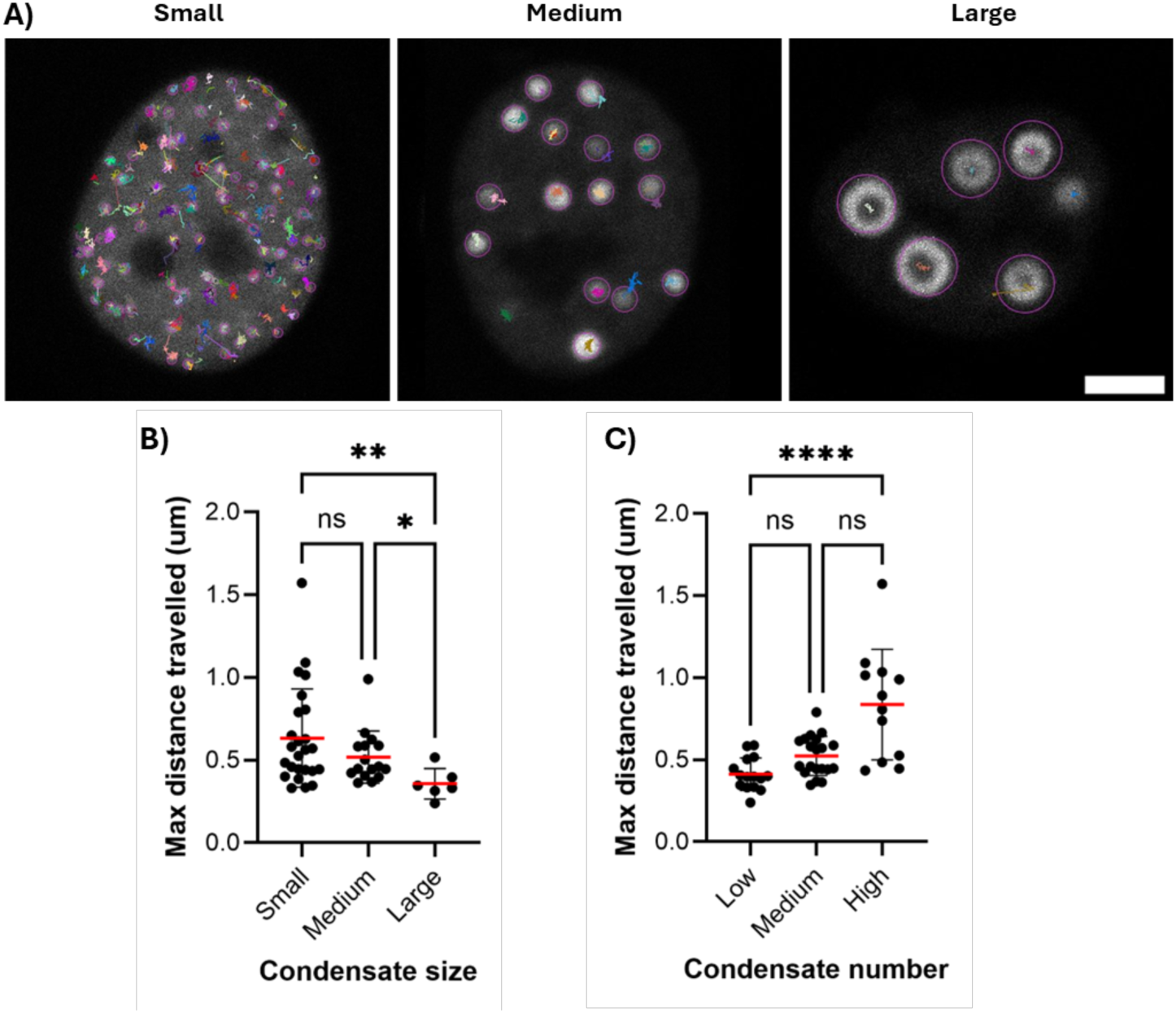
Mobility of ESR1-DAB2 condensates is inversely correlated with their size. **(A)** Representative live-cell confocal images displaying the trajectories of individual ESR1-DAB2 condensates, generated using TrackMate. Panels illustrate the diverse mobility profiles and displacement paths (overlay lines) of individual assemblies across distinct cellular environments, ranging from cells with many small puncta to those containing large condensates. The condensates were classified into three groups based on their area: small < 0.5µm^2^, medium 0.5-1.5µm^2^, and large assembly size >1.5µm^2^. Scale bars = 5µm across all panels. **(B-C)** Quantitative analysis of ESR1-DAB2 condensate mobility profiles. **(B)** Maximum distance vs. condensate size: Mean maximum distance travelled per cell, with cells categorised into groups based on their mean condensate size (small < 0.5µm^2^, medium 0.5-1.5µm^2^, and large assembly size >1.5µm^2^; Kruskal-Wallis test, p=0.0087). **(C)** Maximum distance vs. condensate number: Mean maximum distance travelled per cell relative to total number of condensates, categorised groups based on the number of condensates per cell (low <20, medium 20-45, high > 45; Kruskal-Wallis test, p<0.0001). Data were collected from three independent biological repeats. For condensate tracking analysis (panels B-C), data were analysed across distinct cohorts based on condensate area (n = 25 small, n = 16 medium, and n = 6 large assemblies) and condensate number (n = 15 low, n = 20 medium, and n = 12 high).

### Condensate growth restricts the spatial mobility and molecular exchange of *ESR1* fusion proteins

To determine how the segregation of *ESR1* fusion proteins into phase-separated compartments alters their mobility we utilised Slimfield single-molecule tracking (25) to measure their diffusion coefficients (*D*) within the nucleoplasm (26). This revealed a progressive decline in molecular mobility as condensates increase in size (Figure 6A). More specifically, the mean nucleoplasmic diffusion coefficient dropped by nearly 30% in cells containing medium (mean *D* ± SD = 0.44µm^2^/sec ± 0.1; p = 0.0055) and large condensates (mean *D* ± SD = 0.45µm^2^/sec ± 0.07; p = 0.001), compared to cells displaying a diffuse expression profile (mean *D* ± SD = 0.63µm^2^/sec ± 0.09), while cells with small puncta maintained an intermediate mobility (mean *D* ± SD = 0.57µm^2^/sec ± 0.11). This trend indicates that condensate growth restricts the mobility of *ESR1* fusion proteins, which is likely driven by higher molecular crowding or structural anchoring within the condensate network.

**Figure 6:**
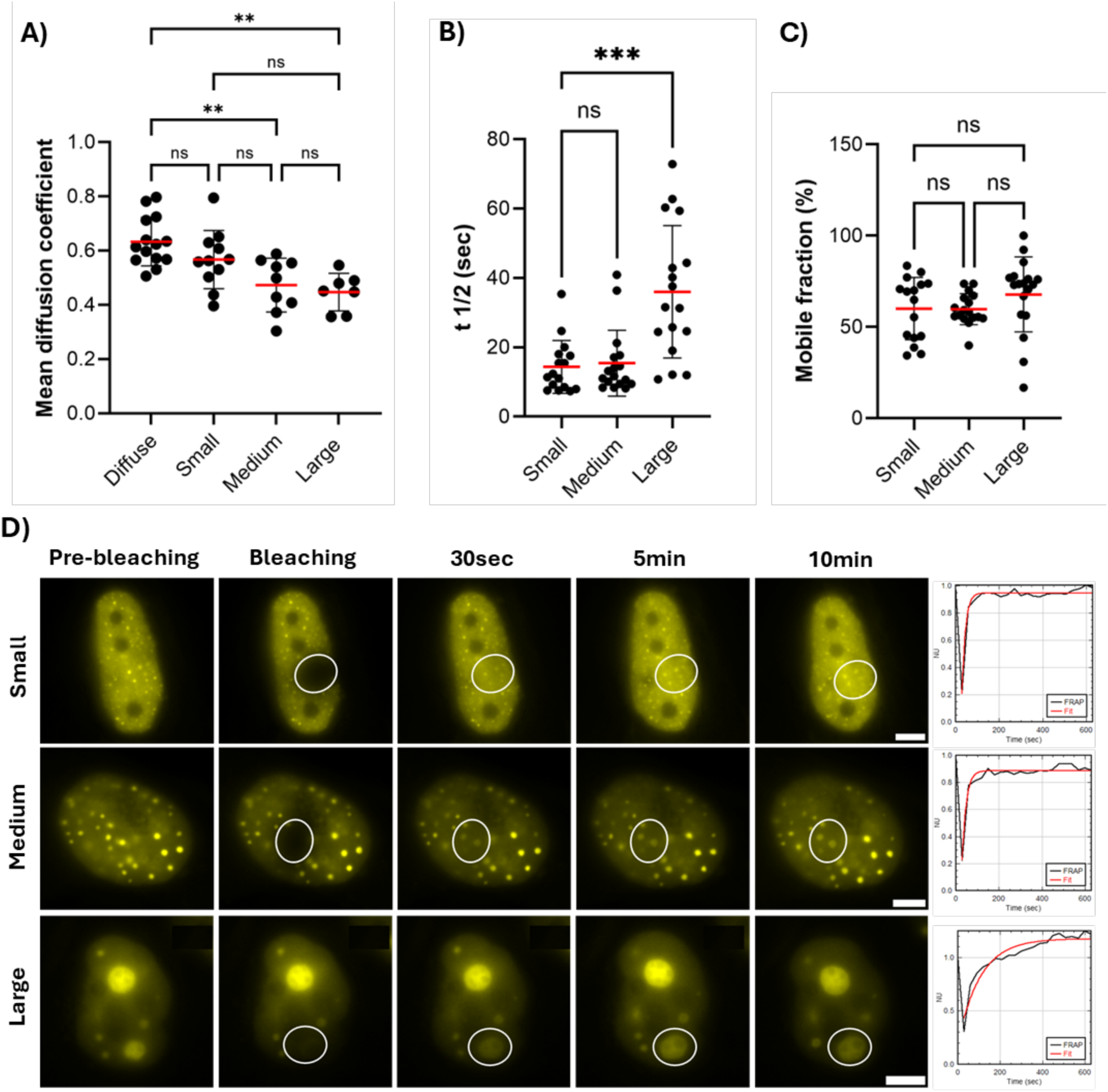
ESR1 fusion protein mobility and molecular exchange. Condensates were categorised into three size cohorts: small <0.5µm^2^, medium 0.5-1.5µm^2^ and large >1.5µm^2^. **(A)** Slimfield single molecule tracking of ESR1-DAB2. The diffusion coefficient (D) of ESR1-DAB2 fusion proteins was measured across diffuse and phase-separated cell cohorts (Kruska-Wallis test, p = 0.0003). Data were collected from three independent biological repeats, with a total of n = 41 cells analysed (n = 14 diffused, n = 11 small, n = 9 medium, n = 7 large). **(B-D)** Fluorescence recovery after photobleaching (FRAP) experiments of ESR1-DAB2 assemblies. **(B)** Half-life (t1/2) across size cohorts (Kruskal–Wallis test, p < 0.0002). **(C)** Mobile fraction (%) representing the extent of fluorescence recovery across size cohorts (Kruskal– Wallis test, p = 0.0725). **(D)** Representative confocal images capturing the fluorescence recovery kinetics of ESR1-DAB2 condensates of various sizes. Panels show individual cells at key experimental intervals: pre-bleaching, initial photobleaching (t = 0), and progressive recovery phases (t= 30s, t= 5min, and t = 10min). Scale bars = 5µm across all panels. Accompanying traces display representative normalised FRAP recovery curves plotting arbitrary units (AU) of fluorescence intensity over time (sec) to illustrate the distinct recovery profiles of each size class. Data were collected from three independent biological repeats, with a total of n = 50 cells being analysed (n = 16 small, n = 17 medium, and n = 17 large assemblies). All graphs display individual cellular data points with lines representing the mean ± SD. Statistical significance evaluated using the Kruskal-Wallis test, followed by Dunn’s post-hoc multiple comparisons test between individual groups.

To evaluate further the molecular mobility within condensates and at the condensate boundary, we performed Fluorescence Recovery After Photobleaching (FRAP) on *ESR1* fusion condensates of varying sizes. Notably, due to technical limitations, we could not bleach the smaller puncta individually; instead, we bleached a localised cluster of these smaller assemblies. Condensates of all sizes showed measurable fluorescence recovery, confirming protein turnover; however, the extent of recovery varied with condensate area with smaller condensates recovering significantly faster than larger ones (Figure 6B-D). Small (<0.5µm*^2^*) and medium (0.5-1.5µm*^2^*) assemblies maintained rapid recovery kinetics (t1/2 = 14.34sec and 15.42sec, respectively), whereas large assemblies (>1.5µm*^2^*) showed a more than two-fold increase in recovery time, averaging a t1/2 of 34.40 seconds (p = 0.0010, Figure 6B). Conversely, the mobile fraction remained stable across the entire population, averaging 62.74% (p = 0.073, Figure 6C). Notably, the greater variability in mobile fraction values observed likely reflects the technical challenges of tracking highly mobile structures that move across focal planes during imaging.

Together, these results indicate the growth of large assemblies simultaneously slows down free diffusion throughout the nucleus and reduces the rate of molecular turnover at the condensate boundary. This strengthens the evidence that larger condensates undergo gel-like maturation or are characterised by a higher internal viscosity that limits the speed of molecular exchange. Nevertheless, the substantial mobile fraction observed across all sizes confirms that while maturation may limit the rate of exchange, larger assemblies still retain a degree of material plasticity and do not become completely impermeable.

### *ESR1* fusion condensates exhibit size-dependent reversibility upon chemical perturbation

To assess whether *ESR1* fusion condensates are reversible, consistent with the LLPS model, transfected cells were treated with 1,6-hexanediol (HD), an aliphatic alcohol widely used to disrupt the weak hydrophobic interactions that stabilise liquid-like condensates (24, 27). Treatment with 2.5% 1,6-HD led to the rapid dissolution of smaller and intermediate condensates, resulting in a diffuse nucleoplasmic distribution of the fusion proteins (Figure 7A). In contrast, larger assemblies remained partially resistant to dissolution, exhibiting only a modest reduction in size. This observation aligns with our single-molecule tracking and FRAP kinetics, reinforcing the idea that larger condensates mature over time into a more resilient, gel-like state. Notably, increasing the HD concentration to 5% or 10% did not enhance the dissolution of these larger assemblies but instead induced significant cellular toxicity and cell death (Supplementary figure 2A,B).

**Figure 7:**
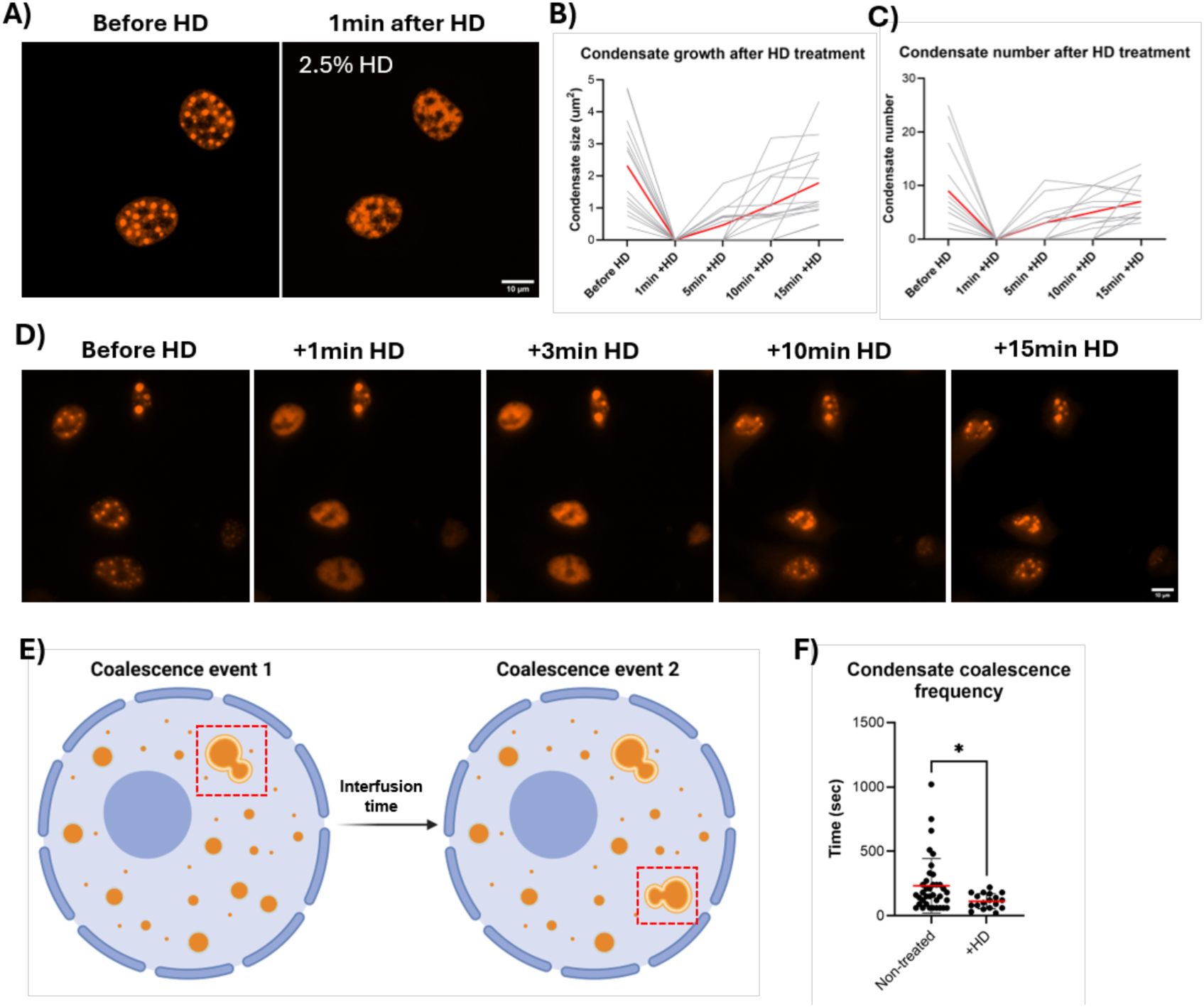
Dissolution of ESR1-DAB2 condensates upon 1,6-hexanediol treatment. **(A)** Representative confocal live-cell images of cells containing ESR1-DAB2 condensates, before and after treatment with 1,6-hexanediol (HD) at final concentrations of 2.5%. Scale bar = 10µm across all panels. **(B-C)** Quantification of condensate growth kinetics over time following complete dissolution with 2.5% HD. Individual cellular trajectories are represented by the grey lines; red lines indicate the cohort mean (n = 15 cells). **(D)** Representative widefield time-lapse images of cells containing ESR1-DAB2 condensates, showcasing the dissolution and growth of assemblies over time after 2.5% HD treatment. Scale bar = 10µm across all panels. **(E)** Schematic illustrating the calculation of the interfusion interval, defined as the time elapsed between consecutive coalescence events within a cell. **(F)** Coalescence frequency of ESR1-DAB2 condensates in unperturbed cells compared to recovering cohorts after 2.5% HD treatment. Data show the interfusion time, defined as the duration between consecutive fusion events calculated per condensate (Mann–Whitney U test, p = 0.0142) The graph displays data from independent coalescence events pooled from multiple cells (n = 14 cells in untreated group; n = 12 cells in treated group) with lines representing the mean ± SD.

Interestingly, long-term live-cell imaging revealed that the dissolved condensates gradually re-formed over time, even under continuous exposure to HD (Figure 7B-D). For the condensates that underwent complete dissolution, the recovery process was remarkably rapid with assemblies nucleating within five minutes and subsequently growing and fusing to restore the pre-dissolution phenotype (Figure 7B-C). The interfusion interval was subsequently calculated by measuring the time elapsed between consecutive coalescence events (Figure 7E). The interfusion interval of these re-forming condensates was significantly lower than that of unperturbed assemblies (time ± SD = from 232.7 ± 211.9s in unperturbed conditions to 112.8 ± 58.59s during reassembly; p = 0.0142; Figure 7F), indicating that the high local concentration of the fusion protein at the site of dissolution lowers the physical barrier for reassembly. Furthermore, condensates consistently reappeared at the same nuclear locations as before dissolution (Figure 7D), highlighting that *ESR1* fusion assemblies are not stochastically distributed. Instead, it indicates that the nuclear environment may contain pre-defined, highly permissive genomic regions that act as stable nucleating scaffolds to direct the localisation of condensate assembly.

### Chromatin accessibility modulates the nucleation and growth of *ESR1* fusion condensates

To investigate the spatial relationship between *ESR1* fusion condensates and the local nuclear environment in more detail, we performed structured illumination microscopy (SIM) following DNA staining. High-resolution imaging revealed that *ESR1* fusion condensates preferentially localised to low-density chromatin regions, while often found adjacent to or encapsulated by dense heterochromatin pockets (Figure 8A). To quantify this spatial distribution, we generated radial fluorescence intensity profiles of Hoechst staining relative to the condensate centre at fixed intervals for a total of 30 pixels (∼1 µm) (Figure 8B). Across all condensate sizes, the Hoechst intensity was the lowest within the core and steadily increased towards the periphery, confirming that the interior of these assemblies is chromatin-depleted. Notably, while small condensates exhibited a sharper, more rapid increase in chromatin density, this intensity gradient became progressively less steep in medium and large assemblies, reflecting the greater distance needed to reach the surrounding heterochromatin boundary. Together, these findings indicate that the assembly of *ESR1* fusions condensates is physically confined by the surrounding chromatin architecture.

**Figure 8:**
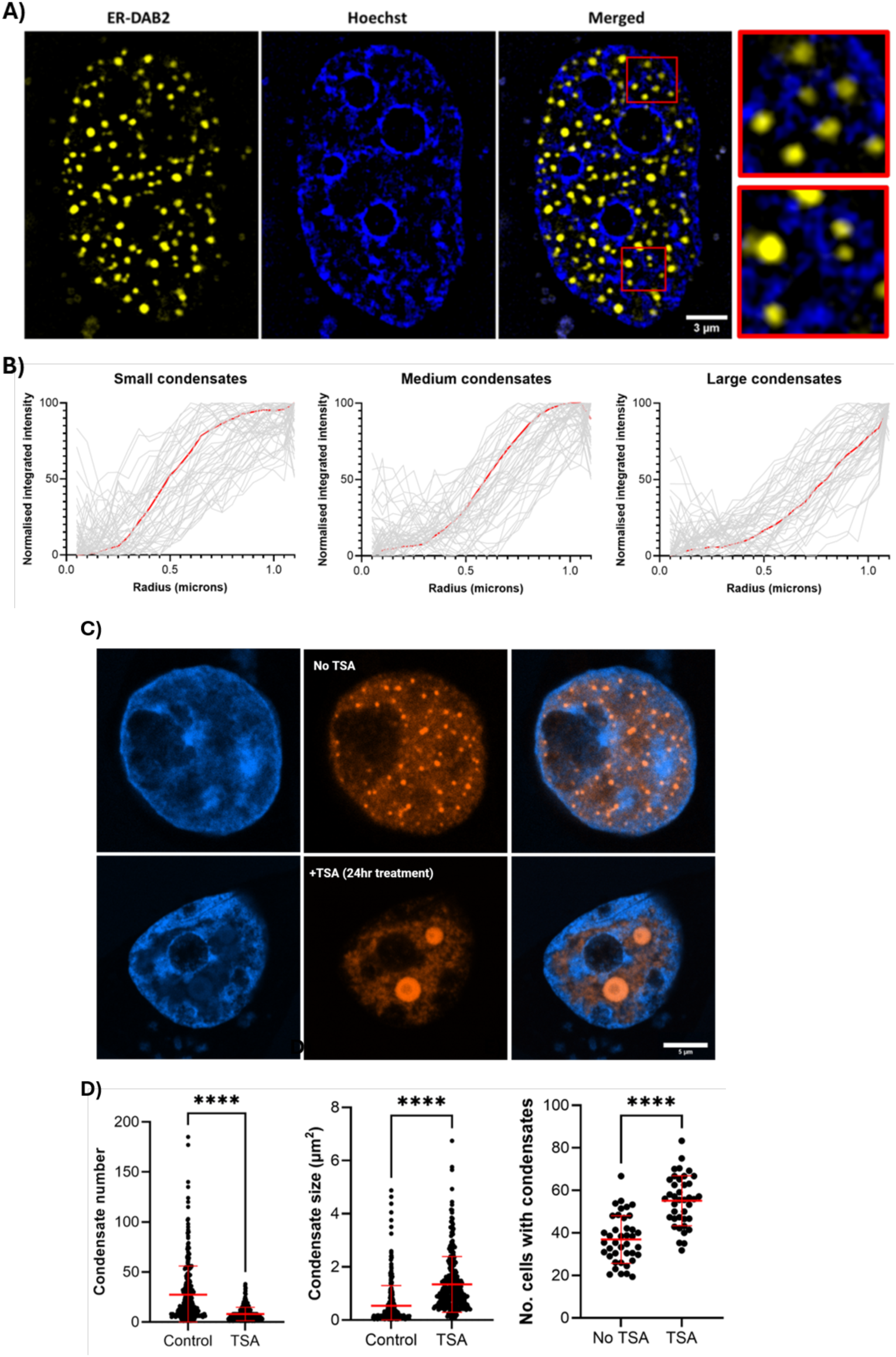
Chromatin architecture modulates the spatial localisation and morphology of ESR1-DAB2 condensates. **(A)** Representative structured Illumination Microscopy (SIM) images showing the localisation of ESR1-DAB2 condensates in relation to chromatin (Hoechst). Merged panels demonstrate the localization of condensates in low-density euchromatin areas, surrounded by dense heterochromatin boundaries. Scale bar = 3um across all panels. **(B)** Quantitative radial fluorescence intensity profiles of Hoechst staining measured from the centre of small (n = 51), medium (n = 53), and large (n = 51) ESR1-DAB2 condensates at fixed intervals for a total of 30 pixels (∼1 µm). The x-axis represents radius (µm) and the y-axis displays the normalized integrated density. Individual profiles are shown in grey, and the red line indicates the mean intensity profile for each size class. **(C)** Representative confocal images of ESR1-DAB2 condensates and DNA, with and without Trichostatin A (TSA) treatment (at 200nM concentration for 24 hours). Images show a shift in the morphology of ESR1-DAB2 condensates after treatment, forming large assemblies when chromatin is relaxed. Scale bar = 5µm across all panels. **(D)** Quantitative analysis of **condensate number** (p<0.0001), **condensate area** (p<0.0001), and **the number of cells containing condensates** (p<0.0001), with and without TSA treatment. Data were collected from three biological replicates (control: n = 305 cells; TSA-treated: n = 310 cells). Graphs display individual cellular data points with lines representing the mean ± SD. Statistical significance evaluated using a Mann-Whitney U test.

To further elucidate the functional impact of the nuclear microenvironment on condensate assembly, cells were treated with the histone deacetylase inhibitor Trichostatin A (TSA) to induce chromatin relaxation (Figure 8C). TSA treatment altered condensate morphology, leading to a significant increase in condensate area (from 0.54 to 1.34μm^2^; p < 0.0001), coupled with a reduction in the total number of puncta per cell (from 27.38 to 8; p < 0.0001) (Figure 8D). Furthermore, the overall proportion of cells containing condensates increased by 49.3% following TSA treatment (p < 0.0001; Figure 8D), indicating that a relaxed chromatin environment is more permissive to the initial nucleation of *ESR1* fusion proteins.

### *ESR1* fusion condensates form elastic, chromatin-constrained networks

To further distinguish between the liquid-like and gel-like nature of *ESR1* fusion condensates, optical trapping experiments were performed to probe the biomechanical behaviour of these assemblies within live cells. The optical trap was moved across nuclei containing condensates to assess their response to applied force. When trapped, condensates exhibited some displacement (mean displacement ± SD = 0.38μm ± 0.39μm) but consistently recoiled back to their original position upon release, indicating a degree of elasticity (Figure 9A). Although condensates were deformable, they maintained a globular structure and did not dissolve under mechanical stress, confirming that larger condensates may have a more gel-like consistency.

**Figure 9:**
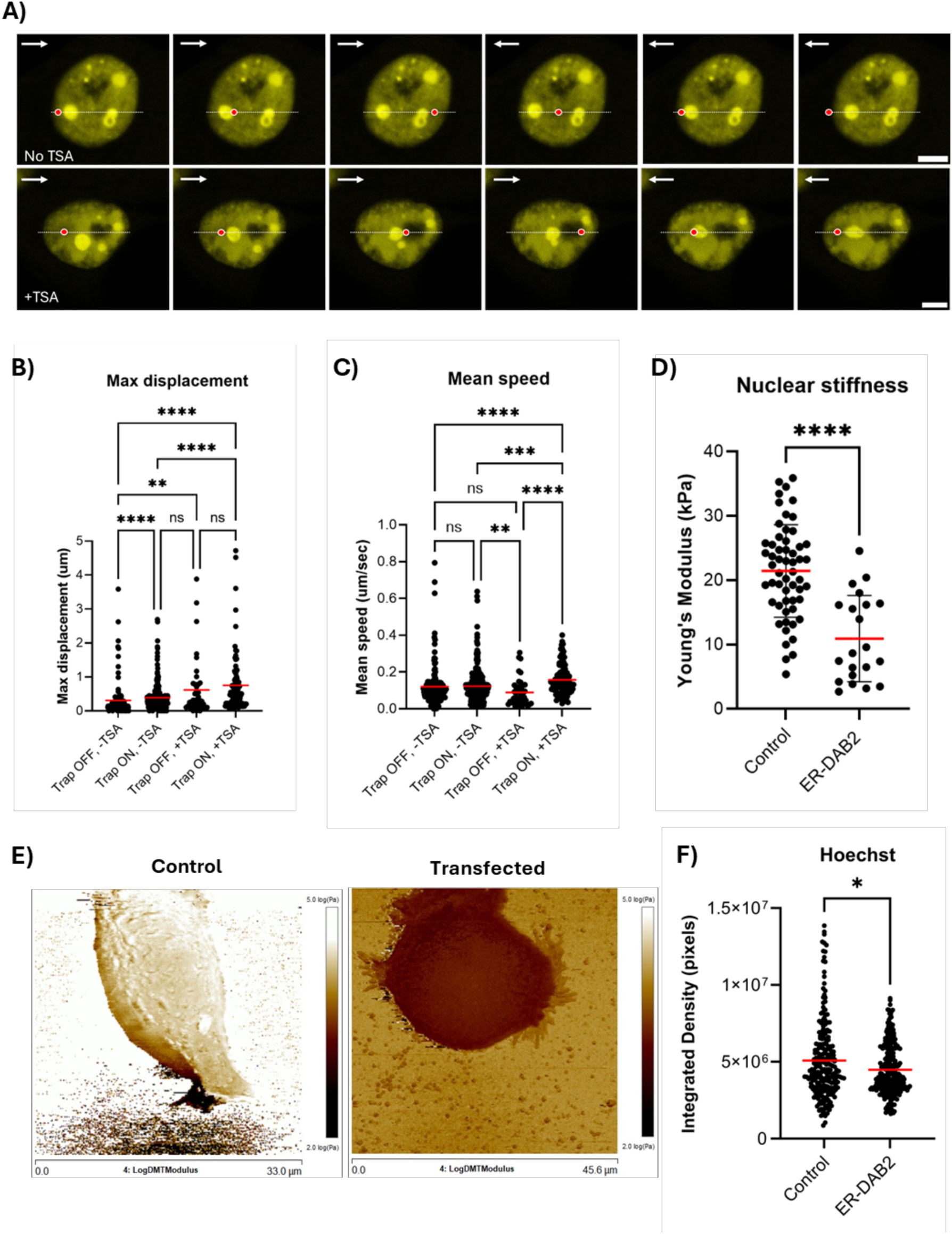
Mechanical properties and mobility of ESR1-DAB2 condensates assessed by optical tweezers experiments. **(A)** Representative confocal images of ESR1-DAB2 condensates upon mechanical perturbation in untreated (–TSA; top panel) and TSA-treated (+TSA; bottom panel) conditions. The white line displays the trajectory of the optical trap, the red dot marks the position of the trap at the time of capture, and the arrow indicates the direction the optical trap is moving. Scale bar = 5µm. **(B-C)** Quantitative analysis of optical tweezer data using TrackMate showing **(B)** maximum displacement and **(C)** maximum speed, of ESR1-DAB2 condensates, before and during trap movement in the presence or absence of TSA treatment. Data were collected from two biological repeats. Graphs display individual condensate data points which represent individual condensates tracked across multiple cells (n = 8 cells for untreated/no trap, n = 14 cells for untreated/+trap, n = 8 cells for TSA-treated/no trap, and n = 10 cells for TSA-treated/+ trap. Solid lines and error bars represent the mean ± SD. **(D)** Quantitative analysis of nuclear stiffness showing the Young’s modulus measurements in control (mock transfection) and ESR1-DAB2 transfected cells (Mann Wittney-U test; p<0.0001). Data were collected from two biological repeats (Control: n = 56 cells; Transfected: n = 21 cells). **(E)** Representative Atomic Force Microscopy (AFM) images showing the Log DMT Modulus for non-transfected (left panel) and transfected (right panel) cells. Graphs display individual cellular data points with lines representing the mean ± SD. **(F)** Quantitative analysis of Hoechst fluorescence intensity as a proxy for DNA density. Data were collected from three biological repeats (Control n = 229 cells; Transfected: n = 260 cells). Graphs display individual cellular data points with lines representing the mean ± SD; Mann Wittney-U test; p= 0.0368.

Additionally, condensates appeared to form an interconnected network, as the displacement of a single condensate resulted in the movement of neighbouring condensates too (Figure 9A, Supplementary video 2B). Collectively, the elastic recoil and coordinated movement suggest that condensates are mechanically constrained within the nuclear environment, likely by chromatin, as observed earlier.

To confirm whether this mechanical constraint was a direct consequence of chromatin compaction, we again utilised TSA-mediated chromatin relaxation. Following TSA treatment, *ESR1* fusion condensates became significantly more deformable and could be displaced over longer distances (mean displacement ± SD = 0.79μm ± 0.85μm compared to 0.38μm ± 0.39μm in untreated controls; p <0.0001; Figure 9B) with reduced recoil to their original position. Moreover, tracking the kinetics of this movement revealed that condensate speed was highest when the optical trap was applied under TSA-treated conditions (mean speed ± SD = 0.16µm/sec ± 0.08 compared to 0.12µm/sec ± 0.08 in untreated controls; p = 0.0006; Figure 9C), indicating that chromatin relaxation significantly lowers the physical resistance to condensate displacement. Despite this increased mobility and speed, condensates retained their structural integrity and did not dissolve. Together, these observations indicate that the formation and dynamics of *ESR1* fusion condensates are heavily influenced by the local nuclear architecture, with chromatin serving as a physical scaffold that limits condensate mobility and dictates the degree of their spatial anchoring.

### *ESR1* fusion condensates alter nuclear architecture

To determine whether *ESR1* fusion condensates can, in turn, influence the global nuclear architecture, atomic force microscopy (AFM) was performed to measure the mechanical properties of nuclei. Force-indentation measurements across the nuclear region revealed a significant reduction in nuclear stiffness in transfected cells relative to control populations (mean Young’s modulus = 21.44 to 10.92 kPa; p < 0.0001) (Figure 9D-E), indicating a shift toward a softer nuclear mechanical state.

To correlate these mechanical changes with structural alterations, we quantified Hoechst fluorescence intensity as a proxy for DNA density. This showed a reduced signal in transfected cells compared to controls (p = 0.0368; Figure 9F), which likely reflects a shift towards a more relaxed, open chromatin state. Together, these findings suggest that *ESR1* fusion condensates do not merely occupy chromatin-depleted regions but also actively remodel the nuclear architecture by promoting chromatin decompaction and altering nuclear mechanics.

## DISCUSSION

### Fusion architecture couples DNA binding with intrinsic disorder to drive the assembly of *ESR1* fusion condensates

Here, we show that several recurrent *ESR1-e6>* fusion proteins form nuclear condensates consistent with LLPS. Condensation appears to be driven by rearrangements that arise at exon 6 of *ESR1*, which eliminate its ligand-binding domain (LBD) while retaining the DNA-binding domain (DBD) and transcription activation function-1 domain (AF-1). This configuration enables fusion proteins to localise to chromatin independently of endocrine signalling. Consistent with this model, we found that the *ESR1* DBD is required for condensate formation, suggesting that chromatin engagement contributes to the nucleation or stabilisation of these assemblies. Another possibility is that deleting the DBD induces broader structural changes or protein misfolding, which may interfere with the interaction surfaces required for self-association. Regardless of the underlying mechanism, these findings demonstrate that the DBD is a critical determinant of *ESR1* fusion protein localisation and spatial organisation.

In addition to chromatin targeting, *ESR1* fusion proteins exhibit increased intrinsic disorder due to partner-derived low-complexity regions. These disordered regions promote the multivalent interactions that drive phase separation. Given that *ESR1* itself contains a disordered transactivation domain (28), fusion partners likely contribute to the interaction capacity rather than acting passively. Consistent with this, many recurrent *ESR1* fusion partners are enriched in intrinsically disordered regions (19), pointing to a distinct structural bias toward configurations that favour higher-order assembly.

Together, these findings support a model in which *ESR1*-exon 6 rearrangements generate fusion proteins that couple chromatin targeting via the *ESR1* DNA-binding domain with enhanced multivalency from partner-derived disorder. Similar principles operate in other oncogenic fusions such as *EWS–FLI1* and *NUP98–HOXA9* where intrinsic disorder and DNA binding cooperate to drive chromatin-associated condensates and aberrant transcription (16, 29, 30). The recurrence of *ESR1-e6>* fusions may therefore reflect selection for architectures that enable phase separation. Defining how variation in fusion architecture and disorder content influences condensate formation and function will be important for understanding *ESR1*-driven oncogenesis.

### *ESR1* fusion condensates exhibit liquid-like properties and undergo size-dependent maturation

*ESR1* fusion condensates display several behaviours consistent with LLPS. Fusion proteins were observed in both diffuse nuclear distributions and discrete puncta, supporting a model in which they nucleate from a soluble pool and grow into larger nuclear assemblies. These condensates showed rounded morphology, fusion upon contact, reversibility following perturbation and dynamic molecular exchange with the surrounding nucleoplasm. Together, these features are consistent with properties reported for transcriptional condensates formed by factors, such as Mediator and BRD4 (22, 31).

Notably, as condensates increase in size, their material properties change. Small condensates are highly dynamic, displaying liquid-like behaviour with rapid molecular exchange and coalescence. As condensates grow, they become less mobile and more viscoelastic, maintaining globular structure under mechanical stress. These observations suggest that condensates retain liquid-like features but acquire viscoelastic properties as they mature. Notably, *ESR1* fusion condensates retain structural integrity without undergoing irreversible hardening as they remain sensitive to 1,6-hexanediol and TSA treatments. This distinguishes them from pathological phase transitions observed in proteins such as FUS, where liquid droplets mature into solid aggregates (32).

The progressive transition from a more liquid-like to a more gel-like state is consistent with the maturation observed in stress granules, nucleoli, and chromatin-associated condensates formed by oncogenic fusions such as *EWS–FLI1* and *NUP98* (29, 32–35). These changes likely arise from increased molecular crowding, strengthened multivalent interactions, or chromatin engagement, raising the possibility that the stabilisation of *ESR1* fusion condensates may prolong their residence at specific genomic loci, thus influencing transcriptional regulation. Nevertheless, it remains unclear whether these assemblies play an active role in regulating transcription or instead represent a biophysical consequence of higher local protein concentration.

### The dynamic interplay between chromatin architecture and *ESR1* fusion condensate formation drives physical confinement and nuclear softening

Chromatin acts as an active regulator of nuclear condensates, shaping their size, mobility, and spatial organisation within the nucleus, while also being remodelled in response to condensate formation (14, 22, 31). Consistent with this reciprocal relationship, our findings indicate that chromatin architecture plays a central role in regulating the assembly and behaviour of *ESR1* fusion condensates. SIM imaging revealed that *ESR1* fusion condensates preferentially localise to low-density, DNA-poor regions bordered by dense heterochromatin. This organisation suggests that condensates form within permissive nuclear pockets, while the surrounding chromatin network acts as a physical constraint that limits their growth, movement and spatial distribution. This is highly consistent with the physical exclusion of chromatin observed for other nuclear condensates, where growing liquid droplets exert mechanical forces that physically push away non-tethered genomic regions, while concentrating targeted loci (36).

Mechanical perturbation experiments further support the idea that *ESR1* fusion condensates are not isolated structures but are instead physically integrated within the nuclear environment. Displacement of a single condensate frequently resulted in the coordinated movement of neighbouring condensates, indicating that these assemblies are mechanically coupled. Such behaviour implies that condensates are constrained by an underlying structural network, likely formed by the surrounding chromatin, rather than behaving as freely diffusing, independent liquid droplets. Pharmacological chromatin relaxation using Trichostatin A (TSA) produced larger, more mobile condensates that diffused further and fused more readily. These findings support a model in which chromatin compaction restricts condensate growth and mobility by acting as a physical boundary within the nucleus.

The observation that condensates repeatedly re-formed at the same nuclear locations following dissolution with 1,6-hexanediol further suggests that condensate positioning is non-random. This spatial recurrence may reflect a form of nuclear spatial memory, in which pre-existing chromatin organisation, local binding sites or structural anchoring regions define preferred sites of condensate nucleation. Alternatively, condensate formation may remodel the local chromatin microenvironment in a way that persists after transient chemical disruption, allowing condensates to re-emerge at previously occupied sites. Distinguishing between these possibilities will require time-resolved analysis of chromatin architecture before, during and after condensate dissolution.

Indeed, our biophysical measurements directly support the capacity of *ESR1* fusion condensates to reshape the nuclear microenvironment. More specifically, AFM revealed reduced nuclear stiffness in cells expressing *ESR1* fusion proteins, indicating global nuclear softening – a phenomenon previously linked to changes in higher-order chromatin organisation (37). This structural shift is further evidenced by our quantitative analysis of Hoechst fluorescence, which showed a lower overall intensity signal and altered spatial organisation characterised by prominent low-density regions. Together, these findings demonstrate a loss of chromatin compaction, suggesting that *ESR1* fusion condensates promote a more transcriptionally permissive nuclear environment by reducing chromatin density and increasing genomic accessibility.

Similar condensate-mediated chromatin remodelling has been reported in other systems. For example, HP1α droplets compact nucleosome arrays and reorganise chromatin architecture (27, 38), while Mediator and BRD4 condensates restructure enhancer topology and concentrate transcriptional machinery at regulatory loci (22, 31). Notably, recent work on the transcription factor FOXA1 showed that its condensates actively dissolve condensed chromatin, increasing local accessibility at FOXA1-bound sites (39). Both the intrinsically disordered regions and the DNA-binding domain were required for this chromatin decompaction, demonstrating a direct mechanistic link between condensate formation and changes in chromatin organisation (39). These findings reinforce our model that *ESR1* fusions are not passive nuclear assemblies, but rather actively modulate the physical state of chromatin.

### Potential roles of *ESR1* fusion condensates in transcriptional regulation and oncogenesis

*ESR1* fusions promote hormone-independent transcription and therapy resistance, but their nuclear mechanism remains unclear. Previous *in vitro* and *in vivo* studies have established that several recurrent fusions including *ESR1-YAP1*, *ESR1-PCDH11X*, *ESR1-SOX9*, *ESR1-ARNT2*, *ESR1-LPP*, *ESR1-NCOA1*, *ESR1-CLINT1*, *ESR1-GRIP1*, *ESR1-TNRC6B*, and *ESR1-DAB2*, confer resistance to estrogen deprivation and selective ER modulators or degraders, such as tamoxifen and fulvestrant (9, 11, 20). Structurally, the loss or disruption of the ligand-binding domain (LBD) in these chimeras leads to constitutive, ligand-independent ER signalling, driving persistent cell growth (9, 11, 40). This sustained signalling triggers the expression of canonical ER target genes, including *GREB1*, *TFF1*, and *PGR* (9, 11), with specific fusions like *ESR1-DAB2* and *ESR1-SOX9* reported to hyperactivate ER transcription by up to 40-fold relative to wild-type ER (11, 20).

Beyond driving therapeutic resistance and proliferation, these fusions actively contribute to the metastatic cascade by promoting a metastasis-associated transcriptional programme. For instance, *ESR1-YAP1* and *ESR1-PCDH11X* remodel the ER cistrome to activate epithelial-to-mesenchymal transition (EMT) genes (9, 11). This includes the induction of the transcription factor SNAI1 (Snail), which represses epithelial targets like E-cadherin and VCAN to enhance cellular motility and metastatic capacity in xenograft models (9, 11). Furthermore, fusions like *ESR1-CCDC170* enhance invasive capacity by upregulating Gab1 – a critical scaffold protein for invadopodia formation – potentially through crosstalk with alternative cytoplasmic signalling pathways (41, 42). However, the mechanism by which these diverse fusions drive such widespread changes is still unknown.

Our findings that *ESR1-e6>* fusions assemble into chromatin-associated condensates may be providing the missing biophysical link, implicating phase separation as a central mechanism contributing to this profound transcriptional hyperactivity. Mechanistically, retention of the DNA-binding domain (DBD) enables chromatin tethering, while their preferential localisation within chromatin-depleted regions may create a permissive environment for hyperactivation. The recurrent preservation of the DBD highlights a strong selection for architectures that couple genomic tethering with the emergent properties of phase-separated condensates, ultimately reinforcing aberrant transcriptional programmes to bypass endocrine therapies. This model is consistent with chromatin-targeted condensates formed by Mediator, BRD4 (22, 31), and oncogenic fusions such as *EWS– FLI1* and *NUP98–HOXA9* (29, 30), which organise transcription at specific genomic loci. *ESR1* fusion condensates may similarly act as persistent hubs that sustain ER signalling independently of estrogen and outcompete wild-type ER, by continuously concentrating the core transcriptional machinery at oncogenic targets.

Beyond their presence at these loci, the functional significance of condensate size and material properties needs to be explored in more detail. Smaller condensates may be highly dynamic and support transient transcriptional bursts but may lack the size or stability required for prolonged regulatory effects. Conversely, very large assemblies could either amplify signalling through enhanced stability and interaction strength or become functionally inert if excessive density restricts molecular exchange. An intermediate condensate size may therefore represent an optimal balance between dynamic exchange and structural persistence (15, 31).

Nevertheless, whether phase separation is strictly required for *ESR1* fusion function, or whether it represents a downstream consequence is an important question that remains to be addressed. Biomolecular assemblies do not always correspond to transcriptional activity, and could alternatively serve other structural or buffering roles, such as modulating protein availability or protecting the fusions from degradation (15), rather than actively driving gene expression. Therefore, defining how condensate assembly, dynamics, and chromatin engagement relate to gene activation will be essential to establish both their mechanistic significance and their potential as therapeutic targets.

## Conclusion

Collectively, our findings define a unifying mechanism by which recurrent *ESR1* fusion events rewire receptor architecture to couple chromatin binding with multivalent interactions, enabling the formation of chromatin-anchored condensates with distinct material properties. Exon 6 rearrangements generate fusion proteins that escape endocrine control, while retaining DNA targeting, driving the assembly of dynamic condensates that mature into stable, chromatin-constrained structures. This coupling between phase separation and chromatin engagement establishes spatially organised nuclear compartments that may stabilise transcriptional outputs independently of estrogen signalling. More broadly, our work positions *ESR1* fusions within a growing class of oncogenic drivers that exploit biophysical principles of phase separation to reprogramme nuclear organisation. Targeting the formation, stability, or chromatin coupling of these condensates may therefore represent a new therapeutic strategy for fusion-driven, endocrine-resistant breast cancer.

## ACKNOWLEDGEMENTS

We thank the UKRI-BBSRC (BB/X008460/1), UKRI-MRC (MR/M020606/1), UKRI-STFC for funding to C.P.T. A.M.G is supported by a studentship from the MRC Discovery Medicine North (DiMeN) Doctoral Training Partnership (MR/W006944/1). A.E.T. is funded by the Wellcome Trust and Royal Society (220192/Z/20/Z). M.C.L. was supported by UKRI-EPSRC (EP/Y000501/1). M.D.N. is supported by a Wellcome Early Career Award (225139/Z/22/Z). The LUMICKS C-trap instrument was funded through MRC Equipment Grant MC PC APP28599 and The Henry Royce Institute for Advanced Materials, funded through EPSRC grants EP/R00661X/1, EP/S019367/1, EP/P02470X/1 and EP/P025285/1.

## AUTHOR CONTRIBUTIONS

**C.P.T.** conceived and supervised the study. **C.P.T.** and **A.M.G.** designed the experiments. **C.P.T.**, **N.F.**, and **A.M.G.** designed and cloned the constructs. **N.F.** and **A.B.** performed the initial characterisation experiments. **A.M.G.** performed the live-cell imaging, SIM, STORM, and AFM measurements with imaging support from **C.P.T.** and **L.W. A.M.G.** performed FRAP measurements with technical guidance and equipment access from **A.E.T. M.D.N.** performed the optical trapping experiments. **A.M.G.** performed Slimfield Microscopy with technical support from **L.F.**, **J.A.L.H.**, and **M.C.L.**, utilising custom Python packages provided by **L.F** for the image processing. All image and quantitative data analysis was conducted by **A.M.G.** with guidance from **C.P.T. A.M.G.** and **C.P.T.** wrote the manuscript, with comments from all authors.

