## Supplementary Data for "Oncogenic ESR1 fusions form chromatin-anchored condensates that remodel nuclear architecture"

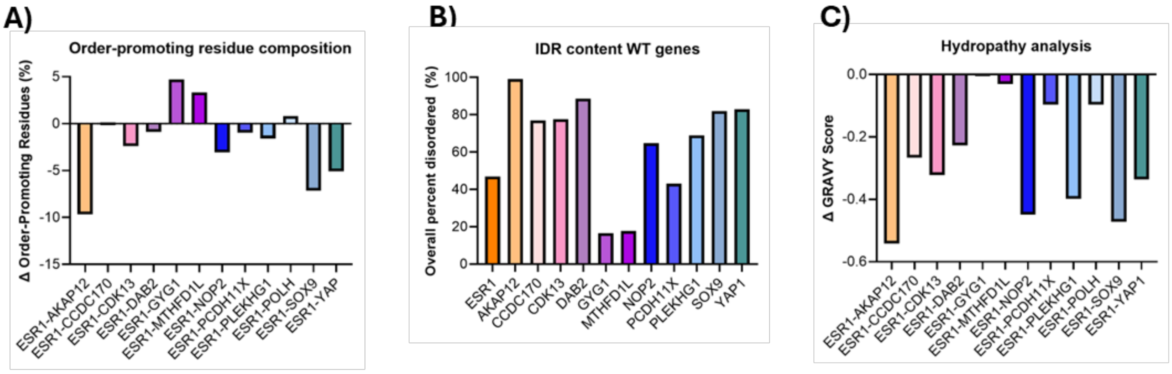


***Supplementary Figure 1: Sequence analysis of ESR1 fusions compared to wild-type ESR1 (WT-ESR1). A)*** *Difference (Δ) in order-promoting residue composition percentage, including tryptophan (W), phenylalanine (F), tyrosine (Y), isoleucine (I), leucine (L), valine (V), cysteine (C), and asparagine (N) which represent bulky hydrophobic and aromatic amino acids. Values were calculated by adding together the percentages of the respective amino acid groups and are displayed as a subtraction difference (Variant % – WT %), where positive values indicate an enrichment in the fusion variant relative to WT-ESR1.* ***B)*** *Total predicted intrinsically disordered region (IDR) content (%) of wild-type gene partners determined using the PONDR VSL2 prediction tool.* ***C)*** *Net difference in Grand Average of Hydropathy (GRAVY) score relative to WT-ESR1 (Variant – WT), calculated using the ExPASy ProtParam prediction tool.*

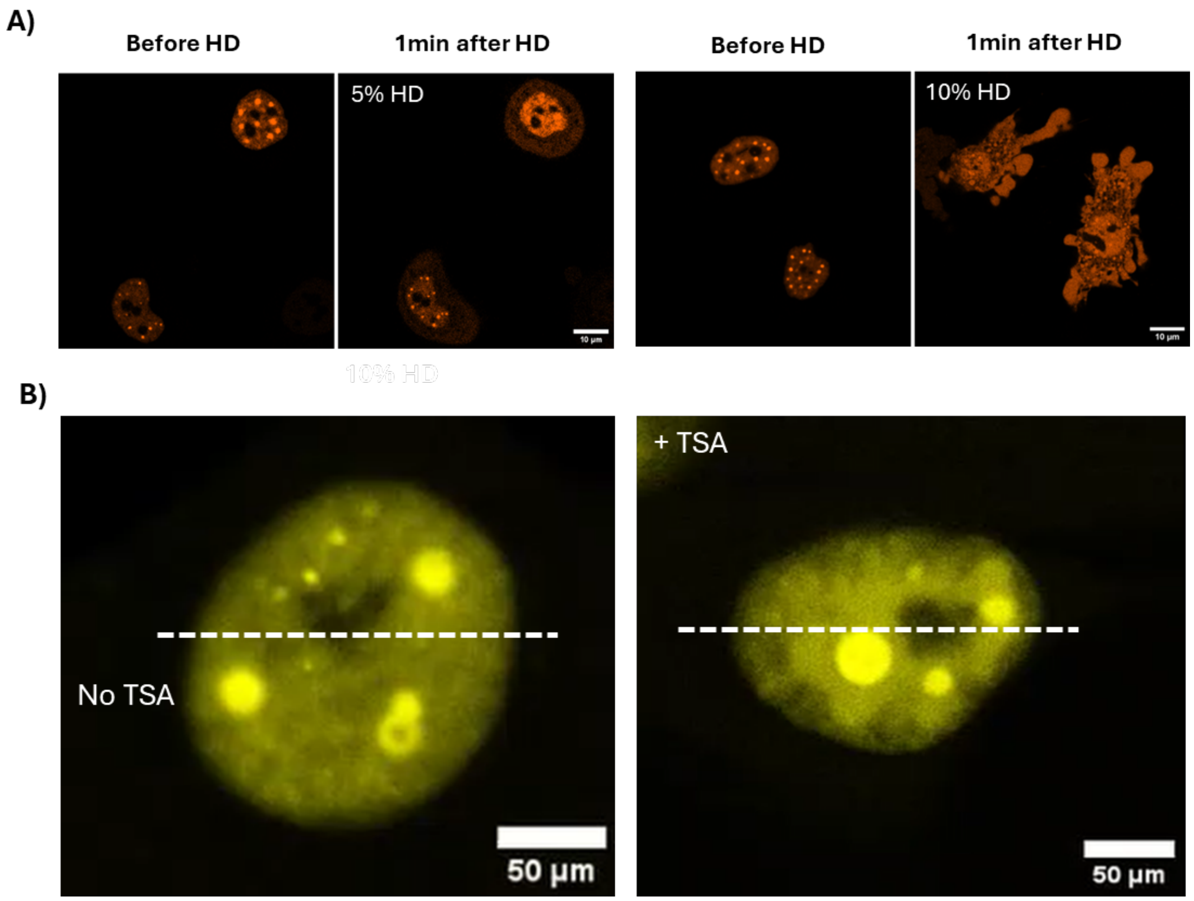


***Supplementary Figure 2: A)*** *Dissolution of ESR1-DAB2 condensates upon 1,6-hexanediol treatment. Representative confocal live-cell images of cells containing ESR1-DAB2 condensates, before and after treatment with 1,6-hexanediol (HD) at final concentrations of 5%, and 10% in the imaging media. Scale bars = 10um across all panels.* ***B) Interconnected condensate network of ESR1-DAB2 condensates.*** *Representative time-lapse videos from an optical tweezers manipulation experiment on ESR1-DAB2 condensates in live cells. The white line traces the trajectory of the optical trap. Movement of a single trapped condensate induces the displacement of neighbouring, untrapped assemblies, demonstrating a highly interconnected network and coordinated viscoelastic mobility. Scale bars = 50µm.*

***Supplementary Table 1: Residue composition and classification of ESR1 fusions compared to WT-ESR1.*** *The table summarizes the total content of hydrophobic, aromatic, positively charged and negatively charged residues of each ESR1 fusion variant and WT-ESR1.*

| *ESR1* variant | Hydrophobic (A, I, L, M, V) | Aromatic (F, W, Y) | Negatively Charged (D, E) | Positively Charged (K, R, H) | Total Charged |
|---|---|---|---|---|---|
| ***WT-ESR1*** | 27.80% | 7.20% | 9.70% | 11.30% | 21.00% |
| ***ESR1-AKAP12*** | 25.30% | 3.10% | 21.20% | 12.40% | 33.60% |
| ***ESR1-CCDC170*** | 33.40% | 4.00% | 15.50% | 17.70% | 33.20% |
| ***ESR1-CDK13*** | 26.60% | 7.10% | 11.00% | 12.10% | 23.10% |
| ***ESR1-DAB2*** | 25.90% | 7.60% | 10.40% | 11.10% | 21.50% |
| ***ESR1-GYG1*** | 30.10% | 9.70% | 10.60% | 12.80% | 23.40% |
| ***ESR1-MTHFD1L*** | 30.70% | 8.00% | 10.60% | 13.30% | 23.90% |
| ***ESR1-NOP2*** | 24.40% | 7.20% | 8.50% | 16.70% | 25.20% |
| ***ESR1-PCDH11X*** | 29.50% | 6.40% | 8.00% | 12.70% | 20.70% |
| ***ESR1-PLEKHG1*** | 25.20% | 6.30% | 12.80% | 13.40% | 26.20% |
| ***ESR1-SOX9*** | 20.40% | 5.40% | 13.00% | 12.20% | 25.20% |
| ***ESR1-YAP1*** | 24.10% | 5.30% | 14.40% | 12.50% | 26.90% |
